# Metastatic infiltration of nervous tissue and periosteal nerve sprouting in multiple myeloma induced bone pain

**DOI:** 10.1101/2022.12.29.522199

**Authors:** Marta Diaz-delCastillo, Oana Palasca, Tim T. Nemler, Didde M Thygesen, Norma A Chávez-Saldaña, Juan A Vázquez-Mora, Lizeth Y Ponce Gomez, Lars Juhl Jensen, Holly Evans, Rebecca E. Andrews, Aritri Mandal, David Neves, Patrick Mehlen, James P Caruso, Patrick M. Dougherty, Theodore J Price, Andrew Chantry, Michelle A Lawson, Thomas L. Andersen, Juan M Jimenez-Andrade, Anne-Marie Heegaard

## Abstract

Multiple myeloma (MM) is a neoplasia of B plasma cells that often induces bone pain. However, the mechanisms underlying myeloma-induced bone pain (MIBP) are mostly unknown. Using a syngeneic MM mouse model, we show that periosteal nerve sprouting of calcitonin-gene related protein (CGRP^+^) and growth associated protein 43 (GAP43^+^) fibres occurs concurrent to the onset of nociception and its blockade provides transient pain relief. MM patient samples also showed increased periosteal innervation. Mechanistically, we investigated MM induced gene expression changes in the dorsal root ganglia (DRG) innervating the MM-bearing bone and found alterations in pathways associated with cell cycle, immune response and neuronal signalling. The MM transcriptional signature was consistent with metastatic MM infiltration to the DRG, a never-before described feature of the disease that we further demonstrated histologically. In the DRG, MM cells caused loss of vascularization and neuronal injury, which may contribute to late-stage MIBP. Interestingly, the transcriptional signature of a MM patient was consistent with MM cell infiltration to the DRG. Overall, our results suggest that MM induces a plethora of peripheral nervous system alterations that may contribute to the failure of current analgesics and suggest neuroprotective drugs as appropriate strategies to treat early onset MIBP.

**Significance statement:** Multiple myeloma is a painful bone marrow cancer that significantly impairs the quality of life of the patients. Analgesic therapies for myeloma-induced bone pain (MIBP) are limited and often ineffective, and the mechanisms of MIBP remain unknown. In this manuscript, we describe cancer-induced periosteal nerve sprouting in a mouse model of MIBP, where we also encounter metastasis to the dorsal root ganglia (DRG), a never-before described feature of the disease. Concomitant to myeloma infiltration, the lumbar DRGs presented blood vessel damage and transcriptional alterations, which may mediate MIBP. Explorative studies on human tissue support our preclinical findings. Understanding the mechanisms of MIBP is crucial to develop targeted analgesic with better efficacy and fewer side effects for this patient population.

## Introduction

Multiple myeloma (MM) is an incurable malignant bone marrow disorder characterized by abnormal immunoglobulemia along with the development of osteolytic bone lesions, hypercalcaemia, renal impairment and anaemia (Kyle and Rajkumar, 2009). Research into the mechanisms of MM has grown exponentially over the last few decades, leading to the introduction of novel therapies such as autologous stem cell transplantation, proteasome inhibitors and immunomodulators as first-line treatment for the disease, which altogether have doubled the median survival time of MM patients (Kumar et al., 2008; Kazandjian and Landgren, 2016). As research attempts to convert MM into a chronic condition, improving the patientś quality of life becomes crucial. In a 2016 systematic review of symptom prevalence across MM patients, pain and fatigue were listed as the most common complaints with over 70% of patients reporting pain, which was described as severe in over 40% (Ramsenthaler et al., 2016). Moreover, a profound disconnect exists between the patientś self-reported pain experience and their physicianś estimation, with recent research suggesting that almost half of attending clinicians underestimate the severity of bone pain in this patient population (Quinn et al., 2022).

Today, pain management in MM patients includes disease-modifying agents targeted to reducing bone disease (i.e. bisphosphonates, denosumab, radiotherapy) and opioids, often in combination with a corticosteroid as adjuvant (Niscola et al., 2010; Coluzzi et al., 2019). On top of well-known side effects of opioids, such as constipation, development of tolerance and risk of addiction, their effect on breakthrough pain, a common occurrence in bone cancer, is limited (Mercadante, 2018). Thus, there is an evident need to unravel the pathogenesis of bone pain in MM, which may lead to mechanism-based strategies to alleviate symptom burden and improve quality of life in a growing patient population.

In a previous study, we established a local, immunocompetent mouse model of myeloma-induced bone pain (MIBP) through intrafemoral transplantation of 5TGM1-GFP cells in C57BL/KaLwRijHsd mice (Diaz-delCastillo et al., 2020b). In this model, we observed the development of pain-like behaviours over time, which were only partially reversed by systemic antiresorptive therapy; moreover, our study showed profound bone marrow denervation at the terminal stages of the model. Here, we elucidate central and peripheral dysregulations driving the onset and maintenance of MIBP.

## Materials and methods

### Cell culture

Mouse 5TGM1-GFP cells passaged *in vivo* were grown in suspension in RPMI media containing glutamine and phenol red, 1% penicillin/streptomycin (100U/100µg/ml), 1% sodium pyruvate (1mM), 1% MEM non-essential amino acids and 10% FBS, at 37°C and 5% CO_2_. All reagents were purchased from Thermo Fischer Scientific, Denmark.

### Animals

Male 5 to 7-weeks-old C57BL/KaLwRijHsd mice from Envigo (Venray, Netherlands) were housed in a temperature-controlled (22 ± 2 °C) room with 50% relative humidity under a 12:12 light:dark cycle (lights on a 07:00 AM). Mice were housed in groups of 4 or 5 in standard individually ventilated GM500^+^ cages (524 cm^2^) in a mouse-dedicated room and allowed to acclimatize >7 days prior to experimental allocation. Cages were enriched with red translucent shelter, an S-brick, paper ropes and corn hidden in the bedding (Tapvei 2HV bedding, Brogaarden, Gentofte, Denmark). Food (Altromin 1314; Brogaarden, Gentofte, Denmark) and water were provided *ad libitum.* Experiments were conducted in accordance with the Danish Act on Animal Experiments (LBK No. 474 of 15/05/2014) and approved by the Danish animal Experiments Inspectorate. This manuscript is reported in accordance to the ARRIVE 2.0 guidelines.

### Experimental design

In a time-course study, mice were intrafemorally inoculated with 5TGM1-GFP myeloma cells or vehicle, their behaviour analysed over time, and euthanized at two different time-points (i.e. post-surgical day 17 and 24, respectively) by transcardial perfusion before tissue collection. In the transcriptomics experiment, DRGs from 5TGM1-GFP or vehicle-bearing mice were collected and fresh frozen for further RNA extraction and transcriptome sequencing 24 days after surgery. In the NP137 experiment, mice were dosed with NP137 (10 mg/kg, i.p.) or vehicle biweekly, their behaviour analysed over time and euthanized at end-point by transcardial perfusion before tissue collection. All behavioural testing was carried out in a quiet room during the light phase (between 07:00 AM and 19:00 PM) by the same researcher, who was blinded to experimental group. Mice were randomized according to baseline burrowing capacity or weight. Good laboratory practices are described in the Supplementary Table S1 (see Supplementary File S1); all materials are available upon request.

### Model induction

Mice were anaesthetized with ketamine/xylazine cocktail (85,5 mg/kg Ketaminol vet-MSD Animal Health, The Netherlands- and 12,5 mg/kg Nerfasin vet – Virbac, Kolding, Denmark; i.p.); eye ointment was applied to prevent dryness (Ophta A/S, Actavis Group, Gentofte, Denmark). Upon confirmation of loss of pedal reflex, the mouse was placed on a heating pad, the leg was shaved and disinfected with 70% ethanol, and an incision (<1 cm) was made above the right anterior knee. The retinaculum tendon was slightly displaced, and the patella ligament pushed to the side, exposing the distal femoral epiphysis, where hole was drilled with a 30G needle. Through an insulin needle (0.3 mL; BD Rowa Technologies, Lyngby, Denmark), 10-µl of vehicle (Hank Balance Salt Solution, Gibco, Denmark) or cell suspension (5x10^4^ 5TGM1-GFP cells) were inoculated into the intramedullary femoral cavity. The hole was closed with bone wax (Mediq Danmark A/S, Brøndy, Denmark), the patella pushed back into place, and the incision closed with surgical clips (Agnthos, Lidingö, Sweden). Mice received 0.5 mL saline (s.c.) and two bolus injections of carprofen (5mg/kg, s.c.; Pfizer, Ballerup, Denmark), one before surgery and another 24 h after.

### Limb use

The limb use of freely walking animals was scored as: 4= normal gait; 3= insignificant limping; 2= significant limping and shift in bodyweight distribution towards the healthy limb; 1= significant limping and motor impairment; 0= paraplegia, as previously described (Diaz-delCastillo et al., 2020b). Briefly, mice were acclimatized with their cagemates to a standard transparent cage of 125 x 266 x 185 mm for ≥10 min. Then, mice were individually transferred to a similar cage where their gate was observed for 3 min.

### Burrowing

Burrowing capacity was assessed as amount of sand (0-3 mm diameter, ScanSand, Herlev, Denmark) displaced from a burrowing apparatus during a 2-hour burrowing session (Sliepen et al., 2019). The burrowing apparatus consisted of a grey opaque plastic tube (200 mm x 72 mm diameter; frontal end raised 30 mm from the ground) filled with 500 g of sand and placed in a transparent plastic cage (125 x 266 x 185 mm) without bedding, closed with a grid lid. Prior to baseline measurements, mice were placed in pairs in the cage containing an empty burrowing apparatus for 2 h. On the second and third day, this procedure was repeated but the burrowing tube was filled with 500 g of sand. Throughout the experiment, burrowing was conducted at the same time and in the same room in absence of the researcher.

### Tissue extraction and analyses

Mice were deeply anaesthetized with a ketamine/xylazine cocktail, as described. Upon loss of pedal reflex, mice were pinned down on their dorsal side and their abdomen and thoracic cage were opened, exposing the beating heart. Transcardial blood was collected with a 27G needle, before transcardially perfusing with 21-28 ml of ice-cold PBS and 4%PFA-0,12% picric acid (Merck, Søborg, Denmark). Spleens were excised and weighed. Ipsilateral femurs were collected, post-fixated 24 h in 4% PFA and stored in 70% ethanol until µCT analyses were performed, and in 0.1M PBS afterwards. Spinal cords and lumbar DRGs were post-fixated 24 h in 4%PFA-0,12% picric acid, dehydrated in 30% sucrose and embedded in OCT (Sakura, Japan). In the transcriptomics experiment, mice were perfused with 10-ml of ice-cold PBS and their lumbar DRGs quickly extracted and fresh-frozen in RNAse-free Eppendorf tubes in a 3-Methylbutanol freezing bath.

### Serum IgG_2b_ analyses

Blood was left undisturbed at RT for 30-90 min and thereafter centrifuged 10 min at 4 °C and 6000 RPM. Serum was stored at -80°C and IgG_2b_ was measured with a sandwich ELISA kit (Bethyl Laboratories, #E99-109; AB_2892024) following manufactureŕs recommendations.

### µCT analyses

Ipsilateral femurs were analysed in a SkyScan 1272 ex vivo scanner at 50 kilovots (kV) and 200 mA, as previously described (Lawson et al., 2015). A 0,5 mm aluminium filter and a pixel size of 4,3 mm^2^ were used. Following standard guidelines (Bouxsein et al., 2010), the following morphometric parameters were assessed: bone volume per total volume (BV/TV), bone surface per bone volume (BS/BV), bone surface per total volume (BS/TV), trabecular thickness (Tb.Th), trabecular spacing (Tb.Sp), trabecular number (Tb.N) and lesion area. Representative 3D models were recreated with ParaViewSoftware (Clifton Park, NY, USA). For the anti-netrin experiments, scans were reconstructed using NRecon software (version 1.6.1.1; SkyScan) within a dynamic range of 0 to 0.15 and a ring artifact reduction factor of 5%. Reconstructed images were analyzed using CTAn (version 1.9.1.1 Bruker, Belgium). Analysis was done on the cross-sectional images of the tibias or femurs at 1.2 mm (offset) from the distal break in the growth plate (reference point), on a fixed 1-mm region. The trabecular bone was carefully traced on all cross-sectional images of the distal femur and BV/TV was calculated.

### Immunohistochemistry in lumbar DRGs

Lumbar DRGs were serially sectioned at 16-µm thickness. Slides were washed thrice, permeabilized in 0.1% Triton X-100/T-PBS and blocked with 1% BSA or 3% NDS before antibody labelling against activating transcription factor 3 (ATF3; either 1:200 polyclonal, rabbit anti-mouse, Novusbio, Abingdon, United Kingdom; #NBP1-85816, AB_11014863, or 1:400 polyclonal, rabbit anti-mouse, Santa Cruz, Dallas, Texas; #SC-188, AB_2258513), tyrosine hydroxylase (TH), a marker for post-ganglionic sympathetic neurons (TH polyclonal rabbit anti-mouse; 1:1000, Millipore; #AB152, AB_390204), CD31, a marker of blood vessel endothelial cells (CD31 monoclonal rat anti-mouse, 1:500, BD Pharmingen, San Diego, CA; #557355, AB_396660), or GFP (GFP polyclonal chicken anti-mouse; 1:2000, Invitrogen; #A10262, AB_2534023). After, slides were washed and incubated 3 h with secondary antibodies: Alexa Fluor 488 (Anti-rabbit, 1:1000, Invitrogen, Themo Fisher Scientific, Slangerup Denmark, #A11008, AB_143165), Cy3 (monoclonal donkey anti-rabbit 1:600; Jackson ImmunoResearch; #711-165-152, AB_2307443), Cy2 (polyclonal donkey anti-chicken, 1:400; #703-225-155, AB_2340370). Sections were counterstained with DAPI (1:10,000, Sigma Aldrich; #D21490) for 5 min, dehydrated in an alcohol gradient and rinsed in xylene. Slides were mounted DPX or fluorescent mounting medium containing DAPI (DAKO Agilent, Glostrup, Denmark).

### DRG neuronal profile quantifications

Serial DRG sections were separated by >120 µm to prevent duplicate counting of neuronal cell bodies. At least a confocal image from three different sections under a 20X objective per animal was taken on a Zeiss LSM 800 confocal microscope (Jena, Germany). The number of ATF3^+^ or TH^+^ immunoreactive neuronal profiles in sensory ganglia were determined in Image J (NIH), as previously reported (Peters et al., 2005), by quantifying both labelled and non-labelled neuronal profiles.

### DRG blood vessel quantifications

Quantification was performed by counting the number of CD31^+^ blood vessels per unit volume (area x thickness) (Ireland et al., 1981; Glaser et al., 2004; Jimenez-Andrade et al., 2008), where blood vessels were identified as CD31^+^ and 2–10 μm in diameter; CD31^+^ branched blood vessels were counted as one single vessel (Weidner et al., 1991).. DRG sections were initially scanned at low magnification (10×) to identify the areas with the highest density of CD31^+^ blood vessels and then a confocal image was obtained at 40× magnification. At least three pictures per animal were taken. An extended depth of focus processing was performed on all Z-stack files for each image and total number and length of CD31^+^ blood vessels within cell body-rich areas (Hirakawa et al., 2004) were determined using Image J (NIH). Then this area was multiplied by the thickness of the section (16 µm).

### Immunohistochemistry in femurs

Following µCT, femurs were decalcified in 10% EDTA for two weeks at 4°C prior to cryoprotection in 30% sucrose and 4°C storage; decalcification was monitored using a portable x-ray equipment (Fona X70, Fona, Assago, Italy). Femurs were longitudinally cut in a frontal plane (25-μm thickness) with a cryostat (Leica 1900, Leica Biosystems, Il, USA), mounted on slides (Superfrostplus, Thermo Scientific; #J1800AMNZ) and allowed to dry (30 min, RT). Subsequently, slides were placed in vertical chambers (Shandon, Sequenza Immunostaining, Fisher Scientific; #73-310-017), washed, blocked (3% normal donkey serum, 0.3% triton X-100 in PBS) for 2 h RT and incubated o/n with a primary antibody cocktail containing anti-GFP antibody (polyclonal chicken anti-GFP, 1:2000; #A10262, AB_2534023; Thermo Fisher Scientific, Rockford, IL, USA), anti-TH antibody (TH polyclonal rabbit anti-mouse; 1:1000, Millipore; #AB152, AB_390204), and an antibody against calcitonin gene-related peptide (CGRP; polyclonal rabbit anti-rat, 1:5000; #C8198, AB_259091; Sigma-Aldrich, St. Louis, MO, USA) or growth associated protein-43 (GAP43; rabbit anti-GAP43; 1:1000; #AB5220, AB_2107282; Millipore, Billerica, MA, USA). Subsequently, sections were washed and then incubated 3 h with a secondary antibodies cocktail of Cy3 donkey anti-rabbit (1:600; Jackson ImmunoResearch; #711-165-152, AB_2307443) and Cy2 donkey anti-chicken (1:400; Jackson ImmunoResearch; #703-225-155, AB_2340370). Slides were counterstained with DAPI for 5 min (1:20,000, Sigma Aldrich; #D21490), washed, dehydrated through an alcohol gradient and xylene, and sealed with DPX mounting medium (Slide mounting medium; Sigma-Aldrich; #06522). *Quantification of the density of nerve fibers on femoral periosteum and neuroma identification* For quantifications of CGRP^+^ and GAP43^+^ nerve fibers density, at least three sections of each mouse femur were analyzed by the same researcher, blinded to model allocation and treatment group. The areas with the highest density of nerve fibers at the metaphyseal periosteum were identified using a 10x objective. In all cases, these areas were located within 0.5 mm-1 mm distance from the distal femoral growth plate. Subsequently, a confocal image was obtained using the Z-stack function from a Carl Zeiss confocal microscope (model LSM 800, Jena, Germany). At least three images were obtained for each marker (20× magnification). The Z-stacked images were analyzed with ImageJ and nerve fibers were manually traced to determine their total length using the freehand line tool. Area of evaluation was manually determined tracing the area of the periosteum. This area was multiplied by the thickness of the section (25 um). Results were reported as the density of nerve fibers (total length) per volume of periosteum (mm^3^) (Mantyh et al., 2010).

For the identification of neuroma-like structures, the following three criteria was followed 1) disordered mass of blind-ending axons (CGRP^+^) that had an interlacing and/or whirling morphology, 2) structure with a size of more than 10 individual axons that is at least 20 μm thick and 70 μm long, and finally 3) a structure that is never observed in the periosteum of normal bone (Devor and Wall, 1976; Sung and Mastri, 1983).

### Evaluation of human bone

Trephine iliac crest bone biopsies from newly diagnosed MM patients were collected at Sheffield Teaching Hospital upon informed consent and under approval from the East Midlands-Derby Research Ethics Committee, UK (REC19/YH/0319). Needle (3 mm) biopsies were collected under local lidocaine anaesthesia by a qualified medical practitioner as routine standard of care. Biopsies were formalin-fixed o/n, decalcified in formic acid, and paraffin embedded before sectioning at 3,5-µm thickness onto DAKO IHC Flex tissue slides (DAKO Aps, Glostrup, Denmark). Sections were deparaffinized in a xylene and alcohol gradient and subjected to antigen retrieval in Tris-EDTA buffer (pH 9.0) o/n. Slides were washed, blocked for endogenous peroxidases and blocked for unspecific staining 20 min in 5% casein/TBS. Sections were labelled with a rabbit anti-PGP9.5 antibody (Sigma Aldrich, Søborg, Denmark; #SAB4503057, AB_10761291), detected with polymeric alkaline phosphatase conjugated anti-rabbit IgG (BrightVision DPVR-AP, Immunologic) and visualized with stay red (DAKO, Glostrup, Denmark). Next, sections were labelled with a mouse mAb against CD34 (Abcam, Cambridge, UK; #ab78165, AB_1566006), detected by polymeric horse radish peroxidase conjugated anti-mouse IgG (BrightVision DPVR-HRP, Immunologic) and visualized with deep space black (DSB, CCC). Sectioned were then blocked with mouse serum and labelled with a mouse mAb against CD138 (BD Pharmnigen, Allschwil, Switzerland; #552723, AB_394443), which was detected with horse radish peroxidase conjugated anti-fluorescein Fab fragments (Merck, Darmstadt, Germany) and visualized with diaminobenzidine (DAB^+^; DAKO, Glostrup, Denmark). Finally, sections were counterstained with Mayeŕs haematoxylin and mounted in Aquatex (Merck, Darmstadt, Germany). Slides were brightfield scanned with a 20x objective in an automated VS200 Slidescanner (Olympus Microscopy, Japan) and analysed in VS200 Desktop, (Olympus) by an experimenter blinded to the slide origin.

### Mouse transcriptomics sequencing and analyses

In the transcriptomics experiment, 20 male C57BL/KaLwRijHsd mice underwent inoculation of 5TGM1-GFP cells or vehicle as described. Pain-like behaviours were assessed over time and mice were euthanized 24 days after surgery by transcardial perfusion with 10 ml of ice-cold PBS. Serum samples were collected and processed for IgG_2b_ assessment as described. Ipsilateral whole DRG lumbar L2-L4 were snap frozen in RNAse free Eppendorf tubes in a 3-Methylbutanol freezing bath. Tissue was homogenized in a bead miller and total RNA was extracted with a Qiagen RNAse microkit (Qiagen, Copenhagen, Denmark). RNA quality was assessed with an Agilent Bioanalyzer and samples with a RIN <8.0 were excluded (n=2).

Remaining samples underwent DNBSEQ transcriptome sequencing by BGI Denmark (BGI, Copenhagen, Denmark). Fastq files with reads with adaptors removed were further used in the bioinformatics analysis upon filtering out low quality ends. We used Salmon (version 1.5.2) (Patro et al., 2017) to quantify transcript expression, using as reference the mouse cDNA set (coding and non-coding transcripts) corresponding to GRCm39. The salmon index was built using the corresponding GRCm39 genome as decoy sequence. Both the cDNA set and genome were obtained from Ensembl, release 104 (Howe et al., 2020; Cunningham et al., 2021). Gene-level expression estimates were further obtained using the tximport R package (Soneson et al., 2015). Genes with non-zero counts in at least 3 samples and with at least 10 estimated mapped reads in at least one of the samples were retained for further analysis. Differential expression analysis between the MM and sham was performed using the Wald test from DESeq2 R package (Love et al., 2014).

Metastatic infiltration of MM cells within the DRG was assessed from the sequencing reads by counting the reads mapping to the eGFP, an artificial transfect to the 5TGM1 cell line. Bbduk (bbmap suite, v. 38.90 (Bushnell) with a kmer size of 31 was used to find the proportion of eGFP matching reads in each sample; eGFP sequences retrieved from (SnapGene).

GSEA (gene set enrichment analysis) for GO biological processes terms and reactome pathways was performed using the R packages clusterProfiler (Sacks et al., 2018) and ReactomePA (Yu and He, 2016). Genes were ranked using the Wald test statistic (stat value) provided by DESeq2. We used the pairwise_termsim() function from enrichplot (Yu, 2019) to obtain the jaccard similarity coefficient (JC) in terms of overlapping gene sets between each two terms. Whenever JC was >0.7, we selected only the term with lowest adjusted p-value value in our enrichment results. Visualizations of the enrichment results were produced using the R packages DOSE (Yu et al., 2015) and enrichplot (Yu, 2019) and data is accessible in GEO under accession code GSE216802.

We compared our MM mouse data with DRG expression data from 6 other mouse models of painful conditions, as described in Bangash et al (Bangash et al., 2018). The transcriptomes of DRGs from the test mice and their corresponding sham had been profiled using the Affymetrix GeneChip Mouse Transcriptome Array 1.0 and obtained from Additional file 3 (Bangash et al., 2018). We compared our model with the 6 models in terms of overlap and direction of differentially expressed genes, as well as in terms of GSEA results. For our MM model we used the set of DEGs defined as padj<0.05, and for the models from Bangash et al, we used the sets of top 300 genes, sorted by pvalue. GSEA analysis for the six models was performed as described for our MM transcriptomic data but ranking the genes by -log(pvalue)*(logFC/abs(logFC). For comparing GSEA results, we selected top 40 most enriched terms in the mouse MM model, sorted by absolute normalized enrichment score, and then extracted the values of the respective terms within the enrichment results corresponding to the other models.

### Human transcriptomics sequencing and analyses

Human DRGs were collected from consenting cancer patients under ethical approval from UT Dallas (UTD) and MD Anderson Cancer Centre Institutional Review Boards. Tissue was extracted during thoracic vertebrectomy in patients presenting malignant spinal tumours; sequencing data from these patients has been previously published by Ray PR et al., where the demographic and clinical characteristics of the cohort are described (Ray et al., 2022). Briefly, DRGs were extracted during spinal nerve root ligation and immediately transferred to cold sterile solution, cleaned and stored in dry ice until sequencing. RNA was extracted using TRIzol^TM^ and cDNA was generated using an Illumina Tru-seq library preparation. Sequenced reads were then trimmed and same-length libraries (38bp) mapped to the GENCODE reference transcriptome (Frankish et al., 2019), as previously described (Ray et al., 2022). Data were obtained from two DRGs belonging to one MM patient and 68 DRGs obtained from 39 patients; further information is compiled in Supplementary file S2. To evaluate gene expression profiles consistent with MM cell infiltration to the DRG, we first created a custom-made marker set from publicly available data. For this, we combined three high throughput datasets of MM cell gene expression based on different technologies: Affymetrix (Barwick et al., 2021), bulk RNA-Seq (Zhan et al., 2006) and single cell RNA-Seq (Jang et al., 2019), and one dataset measuring expression in normal human DRGs (Ray et al., 2018). For the Affymetrix (Barwick et al., 2021) dataset, we obtained the processed data from the GEO NCBI portal using the GeoQuery R package (v.2.60.0) (Davis and Meltzer, 2007). We mapped the probesets of the HGU133Plus2 chip to Ensembl genes using the custom annotation provided by BrainArray (Dai et al., 2005). The file mapping probesets to Ensembl genes was obtained from the BrainArray download page, version 25 (brainarray.mbni.med.umich.edu). Probesets mapping to multiple genes were excluded and when multiple probesets corresponded to the same gene, the one with the highest mean average signal was selected. Probeset intensities were averaged across the 559 replicates, and in total, 16,554 Ensembl genes (of which 15,994 having the biotype “protein coding genes”) were uniquely mapped to probesets on the chip. We assigned ranks to genes by sorting them by decreasing average signal intensity. For the RNA-Seq dataset (Zhan et al., 2006), we used the supplementary table with FPKM counts provided in the Gene Expression Omnibus (GEO) platform (study accession number GSE167968). We averaged expression levels across the 33 replicates and ranked genes by decreasing average FPKM units. For the single cell dataset (Jang et al., 2019), we used the supplementary file with transcripts per million (TPM) counts provided in GEO (GSE118900), and counted the number of cells (out of a total 597) in which each gene has a non-zero expression level. Normal DRG expression levels (Ray et al., 2018) were ranked by the decreasing average TPM counts across the three normal tissue samples. Combining the three MM transcriptomic datasets, we obtained a set of 2528 genes generally expressed in MM cells, i.e. ranked within the top 10,000 genes in both Affymetrix and bulk RNA-Seq and detected in at least 200 of the 597 cells of the single cell experiment. We further reduced this set to a subset of 40 genes by selecting those with an expression rank >15,000 (corresponding to <0.75 TPM) in normal DRG tissue.

We next interrogated MM cell infiltration in human DRG using the MM^D24^ mouse transcriptomic signature as a proxy. For this, we selected the set of upregulated DEGs with a p-adjusted <0.05 in MM^D24^ vs. sham^D24^ mice and evaluated the behavior of their human orthologs in the DRG of MM patients compared to DRGs from patients with other cancer types. For each of the selected markers/DEGs, and for each cancer sample, we computed the percentile expression in relation to the other cancer samples.

### Experimental design and statistical analysis

With exception of the transcriptomics experiments (see above), data were analysed and plotted using Graphpad Prism v.9.3.1 (GraphPad Inc., La Jolla, CA, USA), or SAS 9.4 (SAS Institute, Inc., Cary, NC, USA) and are presented as mean ± standard error of the mean (SEM). Group size was determined in G*Power v3.1.9.7 based on our previously published data (Diaz-delCastillo et al., 2020b) to detect a significant difference in limb use on post-surgical day 25 with a 90% power (α error prob. 0.05). Parametric data were analysed by t-test, one-way ANOVA or repeated measures 2-way ANOVA with Tukeýs correction for multiple comparisons, as required. Non-parametric data was analysed by Friedmańs two-way test followed by Wilcoxońs two-sample test for individual time-points. Transcriptomics data are available in supplementary data or publicly available in GEO (GSE216802); all other data are available upon request.

## Results

### Intrafemoral inoculation of 5TGM1-GFP cells induces nociception

To understand the time course of neuronal changes leading to MIBP and the mechanisms involved, we conducted a time course study where C57BL/KaLwRijHsd mice were transplanted with 5TGM1-GFP MM cells or vehicle into the medullary cavity of the femur and their pain-like behaviours were assayed over time (Figure 1A). A subset of animals was then euthanized on post-surgical day 17 (sham^D17^ and MM^D17^), prior to development of pain-like behaviours (Figure 1B, C) and another on post-surgical day 24 (sham^D24^ and MM^D24^), upon the onset of nociception, as measured by the limb use and burrowing tests (Figure 1D, 1E). The choice of non-stimulus evoked behavioural test was informed by our previous model characterization, where we have demonstrated that this model is not sensitive to mechanical or heat hyperalgesia (Diaz-delCastillo et al., 2019). Locomotor activity test was performed to ensure that the observed behavioural deficits were not a result of impaired motor function (Supplementary Fig S1). Disease development was confirmed by paraproteinemia in both animal cohorts (Figure 1G, I). MM^D24^ mice, but not MM^D17^ mice, also presented splenomegaly (Figure 1F, H), a common feature of 5T MM animal models.

**Figure 1.**
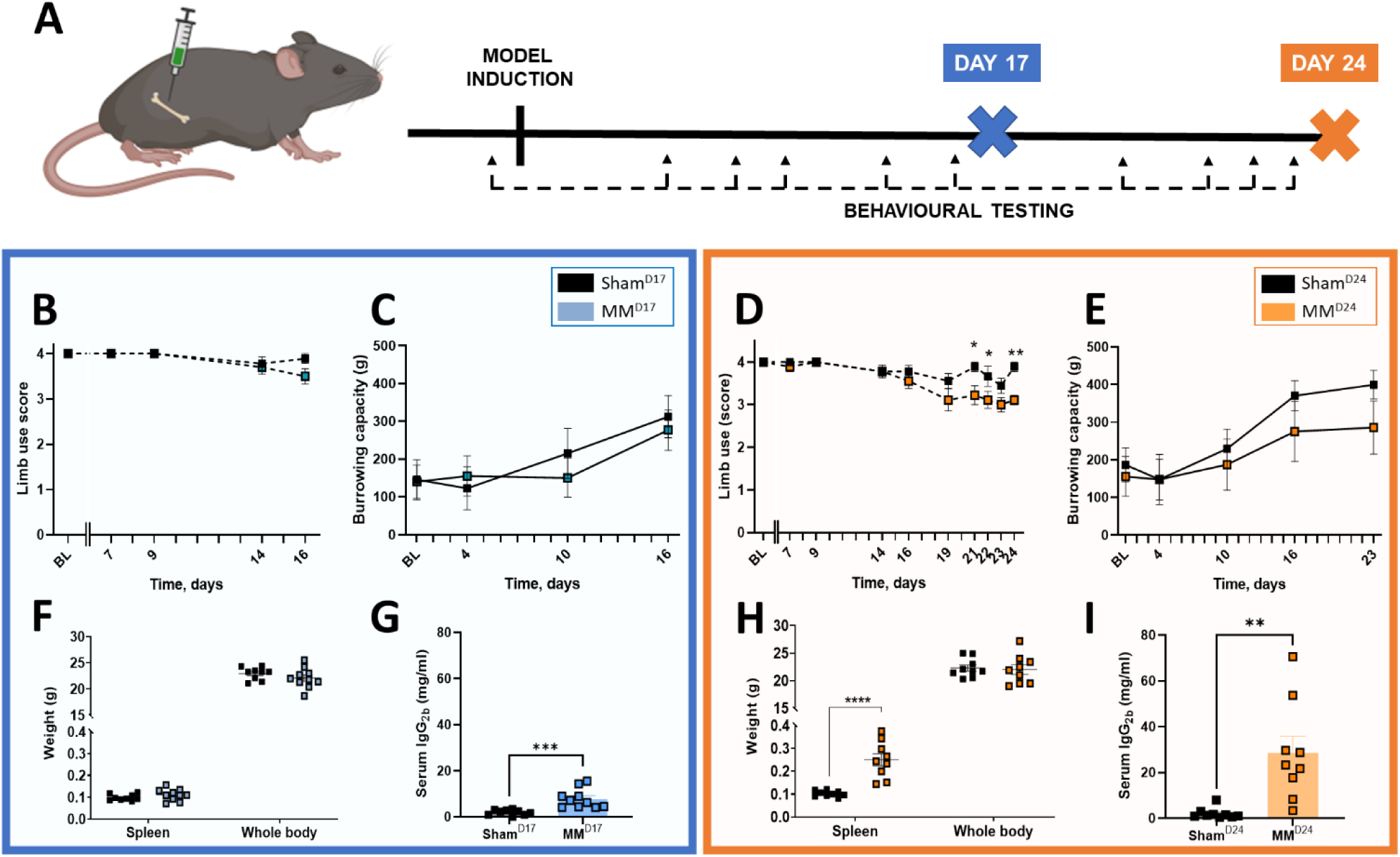
Intrafemoral 5TGM1-GFP cell inoculation in C57BL/KaLwRijHsd mice induces MM disease and pain-like behaviours over time. (A) C57BL/KaLwRijHsd mice were intrafemorally inoculated with 5TMG1-GFP cells or vehicle, their pain-like behaviours were assessed over time, and mice were euthanized on post-surgical day 17 or 24. (B, D) Limb use scores were measured over time. F(1, 180) = 2.10, p=0.0226. Day 21: p=0.0167, Day 22 p=0.0369; Day 24 p=0.0029 by Friedmańs two-way test followed by Wilcoxońs two-sample test; (C, E) Burrowing capacity measured over time. (F, H) End-point spleen and whole-body weight. t(14)=5.98, p<0.0001 by unpaired, two-tailed Student’s *t*-test. (G, I) Endpoint serum levels of IgG_2b_ paraprotein. D17: t(17)=4.068, p=0.0008; D24: t(16)=3.70, p=0.0019 by unpaired, two-tailed Student’s *t*-test. BL= Baseline. Sham n=8-9; MM n=8-10.

### 5TGM1-GFP cells induce a time-dependent pattern of osteolytic lesion development

In previous experiments we observed a partial analgesic effect of bisphosphonates in the localized 5TGM1 model (Diaz-delCastillo et al., 2019), suggesting that MIBP may correlate with osteolysis. Similarly, a plethora of clinical studies have shown that, like in animal models, the clinical analgesic effect of bisphosphonates in MIBP is unclear (Mhaskar et al., 2017). To examine whether the pattern of myeloma-induced osteolysis correlates with the onset of nociception in this model, we performed µCT analyses of the ipsilateral femoral metaphysis of sham and myeloma-bearing bones. We observed that MM^D24^ present a significant increase in cortical lesion area/bone surface (BS), compared with sham^D24^ (Figure 2I, J). In contrast, the lesion area/BS of MM^D17^ remained unchanged, indicating that osteolytic cortical damage develops concurrently to the onset of nociception (Figure 2A, B). Furthermore, our analyses revealed unchanged cortical bone volume in both MM^D17^ and MM^D24^ femurs (Figure 2C, K).

**Figure 2.**
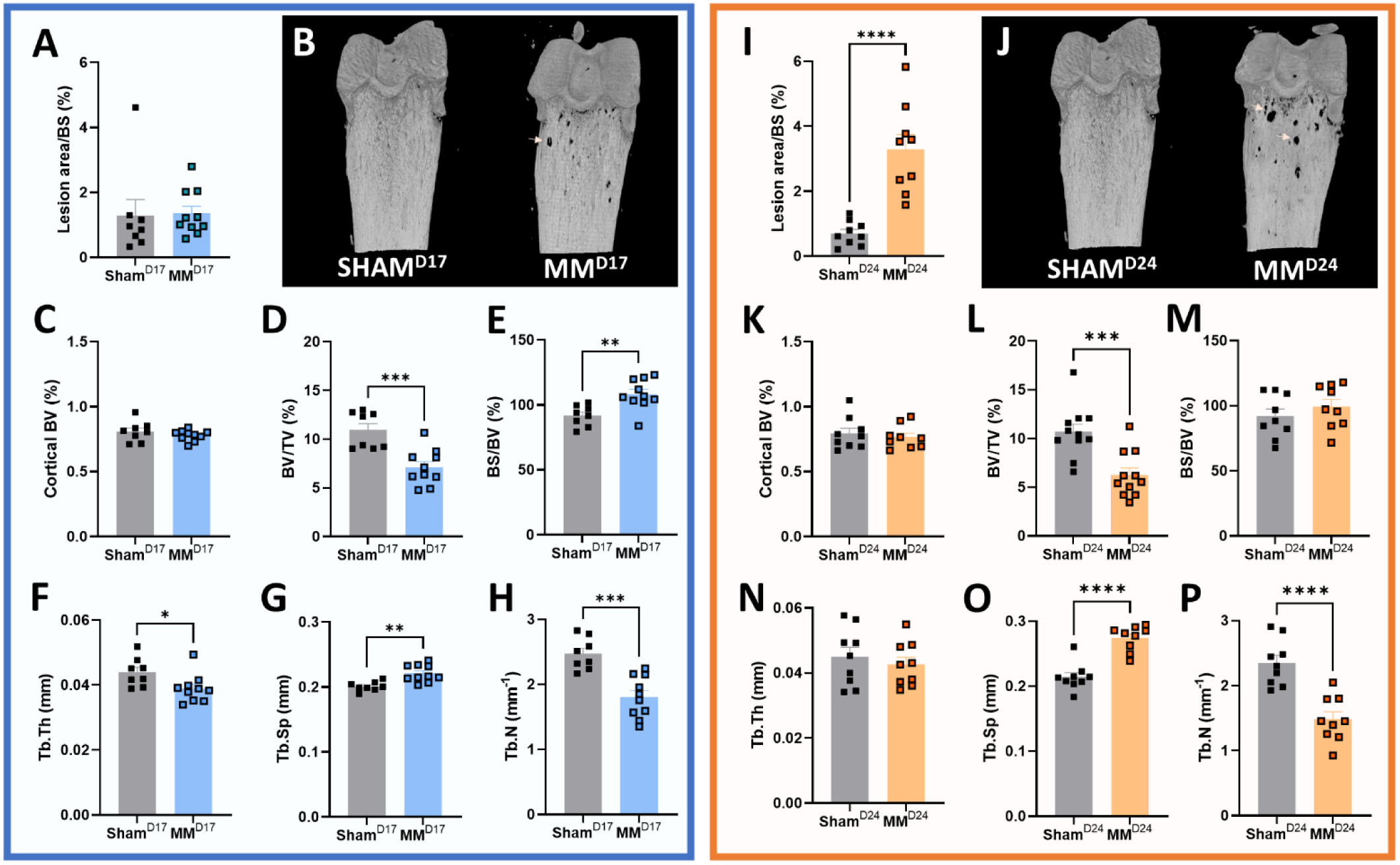
Temporal evolution of myeloma-induced bone disease in 5TGM1-GFP bearing femurs. (A, I) Percentage of lesion area per bone surface (lesion area/BS). t(16)=5.472, p<0.0001 by unpaired, two-tailed Student’s t-test. (B, J) Representative µCT reconstruction images of sham and MM femurs at different time-points; arrows indicate osteolysis. (C, K) Percentage of cortical bone volume (BV). (D, L) Percentage of trabecular bone volume per total volume (BV/TV). Day 17: t(16)=4.323, p=0.0005; Day 24 t(20)=4.107, p=0.0005 by unpaired, two-tailed Student’s t-test. (E, M) Percentage of trabecular bone surface per bone volume (BS/BV). t(16)=3.417, p=0.0035 by unpaired, two-tailed Student’s *t*-test. (F, N) Percentage of trabecular thickness (Tb.Th). t(16)=2.411, p=0.0283 by unpaired, two-tailed Student’s t-test. (G, O) Percentage of trabecular separation (Tb.Sp). Day 17 t(16)=3.903, p=0.0013; Day 24 t(16)=6.381, p<0.0001 by unpaired, two-tailed Student’s t-test. (H, P) Percentage of trabecular number (Tb.N). Day 17 t(16)=4.870, p=0.0002; Day 24 t(16)=5.228, p<0.0001 by unpaired, two-tailed Student’s t-test. Sham n=8-9; MM n=9-10.

Next, we evaluated the effect of 5TGM1 cell inoculation in trabecular bone before and during nociception. Densitometric analyses of MM^D17^ trabecular bone revealed a significant decrease in trabecular thickness (Tb.Th) and number (Tb.N), along with increased trabecular spacing (Tb.S), indicating trabecular bone loss prior to the development of nociception (Figure 2F-H). Moreover, MM^D17^ femurs presented decreased trabecular bone volume per total volume (BV/TV) and increased relative bone surface (BS/BV), compared with sham^D17^ (Figure 2D, E). As expected, MM^D24^ femurs presented a similar pattern of trabecular osteolysis, measured as significantly decreased BV/TV and Tb.N., along with increased Tb.Sp (Figure 2 L-P).

### Multiple myeloma induces periosteal nerve sprouting concomitant to nociception

Our previous studies revealed that 5TGM1 inoculation induced complete bone marrow denervation at the end stages of the model, leading us to speculate that tumour-induced nerve injury contributes to MIBP. To further evaluate the temporal effect of 5TGM1 cell inoculation on the bone marrow microenvironment, we performed immunohistological analyses of sensory (calcitonin-gene related peptide, CGRP^+^) and sympathetic (tyrosine hydroxylase, TH^+^) nerve fibres. We observed that already in MM^D17^ femurs, TH^+^ and CGRP^+^ fibres were not detectable in the bone marrow, which had been colonized by 5TGM1-GFP^+^ cells (Figure 3E, G), suggesting that tumour-induced nerve injury precedes the development of MIBP. To confirm that the absence of marrow innervation was not a result of technical difficulties, we identified both TH^+^ and CGRP^+^ nerve fibres in bones of the sham^D17^ mice (Figure 3B-D) and in the periosteum of sham^D17^ (Fig 3A, C) and MM^D17^ (F, H).

**Figure 3.**
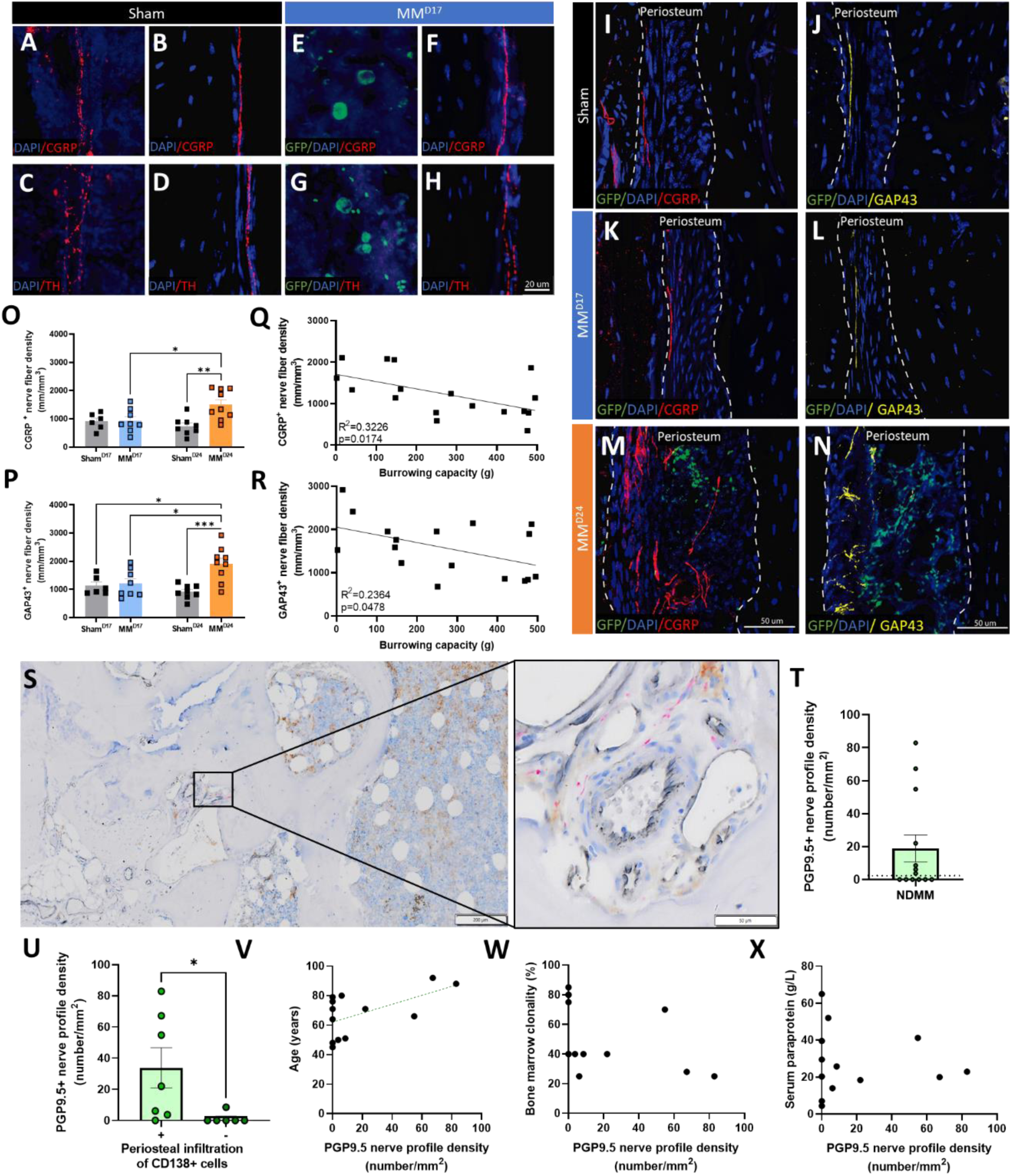
MM induces periosteal nerve sprouting in mouse and human tissue. (A-D) Representative images of CGRP⁺ (A, B) and TH⁺ (C, D) nerve fibres in the bone marrow (A, C) and periosteum (B, D) of sham mice. (E-H) CGRP⁺ (E) and TH⁺ (G) immunoreactivity is undetectable in the bone marrow of MM^D17^, but visible in the periosteal compartment (F, H). (I-N) Representative images of CGRP⁺ (I, K, M) and GAP43⁺ (J, L, N) nerve fibres in the periosteum of sham^D24^ (I, J) and MM^D24^ (M, N). (O) Quantification of relative CGRP⁺ nerve fiber density. F(1,27)=6.343, p= 0.0180; sham^D24^ vs MM^D24^ p=0.0033; MM^D17^ vs MM^D24^ p=0.0354, sham^D17^ vs MM^D24^ p=0.0470 by two-way ANOVA with Tukeýs correction. (R, S) Quantification of relative GAP43⁺ nerve density. F(1, 27)=7.466, p=0.0110; sham^D24^ vs MM^D24^ p=0.0007; MM^D17^ vs MM^D24^ p=0.0206; sham^D17^ vs MM^D24^ p=0.0162 by two-way ANOVA with Tukeýs correction. (Q, R) Correlation between periosteal sprouting and burrowing capacity. CGRP^+^ sprouting: R^2^=0.3226, p=0.0174 and GAP43^+^ sprouting: R^2^=0.2364, p=0.0478 by two-tailed Pearson correlation. (S) Cross-sections of 3.5-µm trephine iliac crest bone biopsies from NDMM patients were triple stained for CD138^+^, CD34^+^ PGP9.5⁺. (T) Quantification of PGP9.5⁺ nerve fibres in the periosteum of NDMM patients; dotted line displays the previously reported human periosteal PGP9.5^+^ nerve density (Sayilekshmy et al., 2019). (U) Periosteal PGP9.5⁺ nerve density in NDMM patients with or without CD138⁺ cell infiltration to the periosteum. t(11)=2.312, p=0.0412 by unpaired, two-tailed Student’s *t*-test. (X-Z) Correlation of PGP9.5^+^ nerve density to age in NDMM. R^2^=0.3305, p=0.0398 by two-tailed Pearson correlation. (X, Z) Lack of correlation between periosteal nerve density and tumour burden measured as bone marrow clonality (Y) or serum paraprotein levels (Z). NDMM= Newly diagnosed multiple myeloma. Sham n= 6-8; MM n=8-9. NDMM n=13.

We next sought to examine the effect of intrafemoral 5TGM1-GFP inoculation on periosteal innervation. The periosteum is the bone compartment with the highest nerve density (Mach et al., 2002; Chartier et al., 2018) and alterations to periosteal nerve fibre innervation have been described as a feature of bone pain (Martin et al., 2007; Mantyh, 2014), including cancer-induced bone pain (Mantyh et al., 2010; Bloom et al., 2011). Our analyses revealed infiltration of MM cells to the femoral periosteum of MM^D24^, which were not present in MM^D17^ femurs. Importantly, we found a significant increase in the density of CGRP^+^ fibres innervating the periosteum of MM^D24^ mice at the onset of nociception, compared with sham^D24^ (Figure 3I, M, O), which was not present at earlier stages (Figure 3K, O). Similarly, the growth associated protein-43 (GAP-34) marker of axonal growth and regeneration demonstrated significant periosteal sprouting in the later stage of MM development (Figure 3J, L, N, P). Altogether, our data suggests that MM cells induce cortical osteolytic lesions that allow escape to the periosteum, where they may promote periosteal nerve sprouting and contribute to the development of nociception. This was further supported by the significant inverse correlation between periosteal CGRP^+^ and GAP43^+^ periosteal nerve sprouting and burrowing capacity (Figure 3Q, R).

To investigate the human relevance of our findings, we next performed an explorative study to evaluate whether periosteal infiltration of MM cells in patients is associated with nerve sprouting. In formalin-fixed, paraffin-embedded trephine iliac crest bone biopsies from 13 newly diagnosed MM (NDMM) patients, we performed a multiplex immunostaining for CD138^+^ MM cells, CD34^+^ blood vessels and the pan-neuronal marker PGP9.5 (Figure 3S). Our quantification showed that the median periosteal nerve density in NDMM patients was 3.736 profiles/mm^2^, ranging from 0 to 82.869 profiles/mm^2^ (Figure 3T); this is in contrast with reports of periosteal nerve density in non-cancerous patients showing a median of 0.077 profiles/mm^2^ (range: 0.02-0.68 profiles/mm^2^) (Sayilekshmy et al., 2019). Moreover, we found a significant increase in periosteal nerve density in NDMM displaying periosteal infiltration of CD138^+^ cells compared with patients without CD138^+^ cells in the periosteum (Fig 3U), suggesting a direct role for MM cells in promoting nerve sprouting. Periosteal nerve density in NDMM patients was positively correlated with age (Figure 3V) but independent of tumour burden, which was assessed as percentage bone marrow clonality (Figure 3W) and paraproteinemia (Figure 3X). Moreover, periosteal nerve density was independent of sex and IgG type (data not shown). This is, to our knowledge, the first evidence of MM-induced alterations to bone innervation in MM patients and altogether our data suggest that periosteal nerve sprouting may play a role in MIBP.

### Pharmacological blockade of periosteal nerve sprouting induces a transient anti-nociceptive effect

Next, we tested the mechanistic role of periosteal nerve sprouting on MIBP using a therapeutic anti-netrin-1 blocking antibody (NP137). Netrin-1 is an axon guidance molecule known to play a pivotal role in neurogenesis through binding to its canonical receptors UNC5 homolog (UNC5H) and deleted in colorectal cancer (DCC) (Madison et al., 2000; Dun and Parkinson, 2017; Boyer and Gupton, 2018). Previous studies have demonstrated a role of netrin-1 on sensory nerve sprouting (Zhu et al., 2019); moreover, silencing netrin-1 reduces hyperalgesia and CGRP+ nerve fiber sprouting in a rat model of disc degeneration (Zheng et al., 2023) and pharmacological netrin-1 inhibition with the NP-137 anti-netrin-1 antibody reduced hyperalgesia in an arthritis model (Rudjito et al., 2021). To investigate whether blockade of periosteal nerve sprouting attenuated MIBP, 5TGM1-GFP inoculated mice were systemically treated with vehicle (MM^VEH^) or NP137 (MM^NP137^; 10 mg/kg, i.p). Biweekly treatment with the anti-netrin-1 antibody did not have an overall behavioural effect, but it delayed the onset of nociception, inducing a significant improvement in limb use scores in MM^NP137^ on day 26, compared with MM^VEH^ (Figure 4A). Moreover, NP137 treatment did not affect overall survival (Figure 4B) nor overall tumour burden, as assessed by terminal splenomegaly (Figure 4C). To evaluate whether the transient analgesic effect was a consequence of decreased osteolysis, we performed µCT analyses of endpoint MM^VEH^ and MM^NP137^ femurs. Our results demonstrated that netrin-1 blockage does not affect BV/TV in MM mice (Figure 4D). Moreover, the structural parameters of trabecular bone and bone mineral density were unchanged (data not shown). To confirm the capacity of NP137 treatment to block periosteal nerve sprouting, we next assessed the presence of CGRP^+^ nerve fibres in the femoral periosteum and found a significant reduction in CGRP^+^ periosteal nerve density in MM^NP137^, compared with MM^VEH^ (Figure 4E-G). Anatomical presence of a microneuroma was observed in 25% of the vehicle-treated MM mice, (Figure 4G), a feature of the disease never before described but that is consistent with animal models of solid bone cancers such as prostate (Jimenez-Andrade et al., 2010) and breast (Bloom et al., 2011) bone metastases or osteosarcoma (Ghilardi et al., 2010; Mantyh et al., 2010). No microneuromas were observed in MM^NP137^ mice. Systemic antibody treatment by itself in sham mice had no effect on behaviour, bone structural parameters or neuronal innervation (data not shown), and pharmacokinetic analyses confirmed the presence of NP137 in the serum of all treated mice (Supplementary Figure S2).

**Figure 4.**
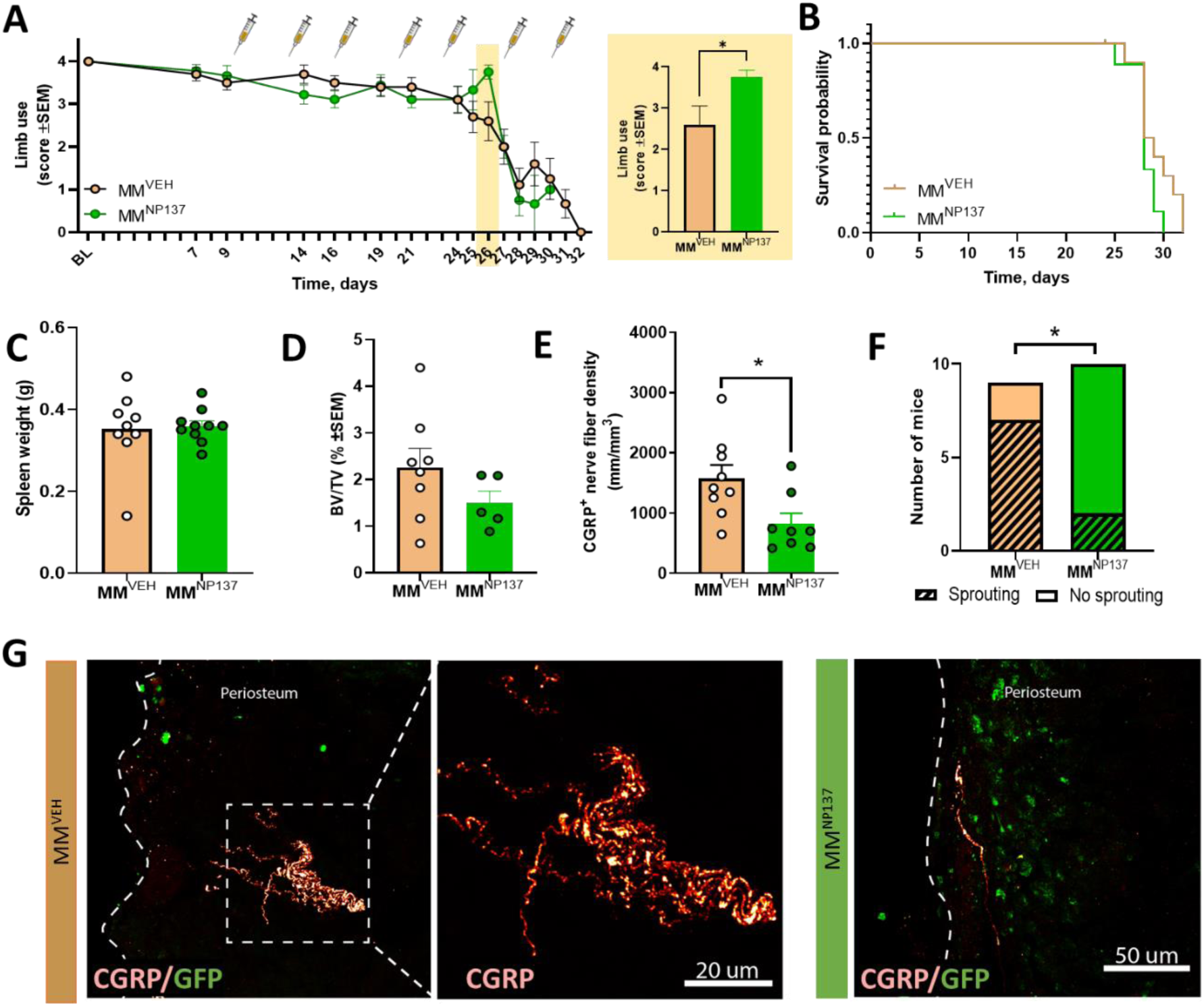
Pharmacological blockage of periosteal nerve sprouting induces a transient anti-nociceptive effect in 5TGM1-bearing mice. (A) Effect of anti-netrin-1 treatment (NP137 10 mg/kg, i.p.; dosing days represented by syringes) or MM mice; exert indicates limb use scores on post-surgical day 26 (onset of pain-like behaviour). t(16)=2.164, p=0.0451 by unpaired, two-tailed student’s t-test. (B) Kaplan-meier curve of vehicle- and NP137-treated MM mice. (C) Endpoint spleen weight. (D) Effect of systemic NP137 treatment on bone osteolysis, measured as bone volume per total volume (BV/TV). (E) Quantification of CGRP^+^ periosteal nerve density. t(15)=2.63, p=0.0188 by unpaired, two-tailed Student’s t-test. (F) Number of vehicle- or NP137-treated MM mice presenting periosteal nerve sprouting. Chi^2^(1)=6.343, p=0.0118 by Chi square test. (G) Representative images of periosteal CGRP^+^ nerve fibres and GFP^+^ MM cells in MM mice treated with vehicle or NP137. MM= Multiple myeloma. MM^VEH^ n=9, MM^NP137^ n=10.

### Dorsal root ganglia (DRG) transcriptomic dysregulation reveals MM infiltration

Following our observation of periosteal sprouting at the onset of nociception, we next hypothesized that MM invasion of the bone niche induces transcriptomic changes in the cell bodies of the innervating nerve fibres. However, considering that blocking of the nerve fibre sprouting only had a transient analgesic effect, there are clearly other mechanisms involved in MIBP. To test this, the DRG transcriptional signature of sham or 5TGM1-GFP inoculated mice during MIBP was evaluated. As expected, MM^D24^ animals displayed nociception (Figure 5A) and splenomegaly (Figure 5B), confirming disease development. On post-surgical day 24, RNA from lumbar DRGs L2, L3 and L4 was extracted and sequenced (if RIN>8), resulting in library sizes of 19-22M reads per sample. The mapping rates to the mouse transcriptome were similar across samples, ranging between 88-92% of all reads. We identified 1.389 differentially expressed genes (DEGs) between MM and sham groups with an FDR<5% (Figure 5C, D). Interestingly, the DRG transcriptomic signature of MM mice was highly heterogeneous across samples; this heterogeneity was not correlated to tumour burden (Figure 5D). However, significant changes in the expression pattern of DRG gene expression were driven by the presence and transcriptional level of green fluorescent protein (GFP) (Figure 5D), suggestive of MM infiltration to the DRG. Next, we performed gene set enrichment analyses (GSEA) to better interpret the transcriptional dysregulation by taking into account the entire set of genes expressed in our data and without setting up any arbitrary threshold of statistical significance on differential expression. GSEA for GO BP (Gene Ontology biological process) terms and Reactome pathways indicated that the main dysregulated signaling pathways in MM mice were related to cell cycle, immune response (activated) and neuronal signaling (suppressed) (Supplementary Figures S3 and S4).

**Figure 5.**
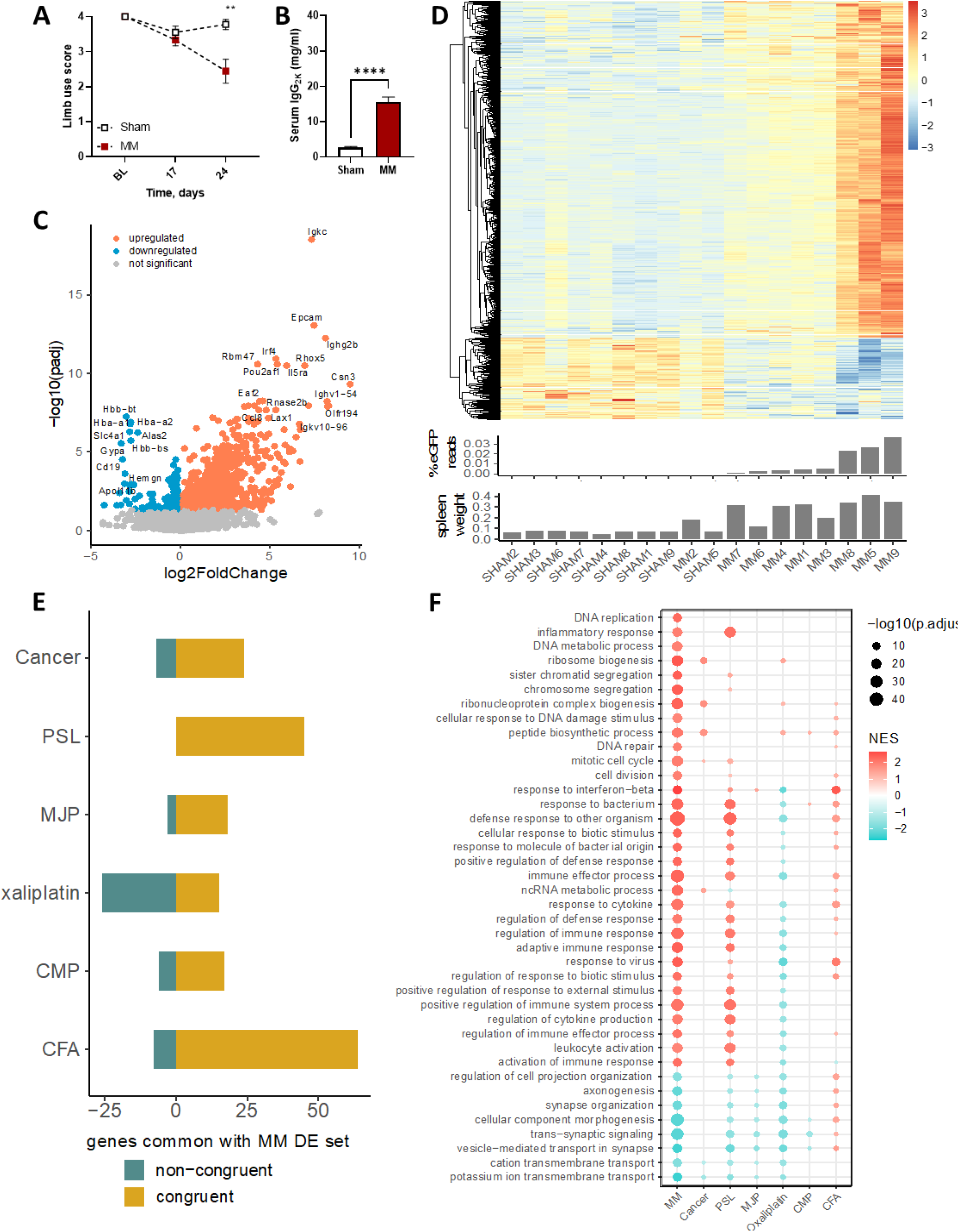
The DRG transcriptional signature in MM unveils MM infiltration to the nervous system and a strong neuropathic component. (A, B) Development of nociception (A) and splenomegaly (B) were confirmed prior to DRG isolation and transcriptomic analyses. F(1, 36) = 0.0127; Day 24 p=0.0012 by Friedmańs two-way test followed by Wilcoxońs two-sample test and t(16)=9.106, p<0.0001 by unpaired, two-tailed Student’s t-test. (C) Volcano plot showing the log2 fold change of differentially expressed genes (DEGs) between sham^D24^ and MM^D24^. (D) Heatmap depicting z-scaled regularized log counts of the 1389 DEGs identified between sham and MM lumbar DRGs (adjusted p-value<0.05). The heterogeneous transcriptome of MM DRG was correlated to the presence and levels of GFP expression. (E) Comparison to other mouse pain models in terms of common DEGs. The overlap between the DEGs identified in MM mice and top 300 DEGs, sorted by p-value, in other painful models, as reported by Bangash et al (Bangash et al., 2018), highlights the neuropathic and inflammatory component of MIBP. In yellow, common DEGs congruent in their direction of regulation between MM and each of the other models; in blue, common DEGs in non-congruent direction of regulation. (F) Model comparison in terms of GSEA. Normalized enrichment scores (NES) and their corresponding p-values for the top 40 enriched GO terms in MIBP are displayed across all seven pain models(Bangash et al., 2018), indicating that the PSL model is the most similar to MIBP. MM= Multiple myeloma. PSL= Partial sciatic nerve ligation. MJP= Mechanical joint loading. CMP= chronic muscle pain. CFA= Complete Freund’s adjuvant. Sham n=9; MM n=9.

Next, we compared the MIBP transcriptomic signature with that of six other models encompassing different painful conditions, as compiled by Bangash et al. (Bangash et al., 2018). These included (Figure 5E) mouse models of painful lung cancer metastasis to the bone (cancer), partial sciatic nerve ligation (PSL), mechanical joint loading (MJL), chemotherapy induced peripheral neuropathy (Oxaliplatin), chronic muscle pain (CMP) and inflammation (Complete Freund’s Adjuvant, CFA). To identify similarities among different painful conditions, we first examined the overlap between the set of MIBP DEGs, and the top 300 genes with smallest p-values from each of the six conditions. Our analyses indicate that MIBP has most DEGs congruent with the PSL and CFA models, suggesting a neuropathic and an inflammatory component (Figure 5E).

Next, we performed (GSEA) for the 6 models and compared results by selecting the top 40 enriched GO BP terms or Reactome pathways in MIBP and visualizing their normalized enrichment score and corresponding adjusted p-values across all models (Figure 5F and Supplementary Figure 4). Interestingly, the transcriptional signature of MM was overall most similar to that of PSL, suggesting a strong neuropathic component in MIBP. These results are in line with our previous finding of periosteal nerve sprouting as a contributing mechanism to MIBP. Even though the overlap of DEGs is higher with the CFA model, the comparative GSEA analysis indicates that this similarity is only retained at the level of common activated inflammation-related pathways. Since spinal microglial reaction is a well-known feature of neuropathic pain (Chen et al., 2018; Inoue and Tsuda, 2018), we characterized the expression of ionized calcium-binding adaptor molecule 1 (Iba1) and phospho-p38 mitogen-activated protein kinase (P-p38 MAPK) in the dorsal horn of the spinal cord of sham^D24^ and MM^D24^ mice. No changes in relative Iba1^+^ or Pp38^+^ cell number were observed at any lumbar region (Supplementary Figure S5). Likewise, no changes in glial fibrillary acidic protein (GFAP) staining were observed, suggesting that astrocytosis is not a main feature of MIBP (Supplementary Figure S5).

### MM infiltration and increased ATF3 expression in DRGs from MM^D24^ mice

To further verify that the GFP reads detected during transcriptome sequencing were caused by MM infiltration to the DRG and not the result of sample contamination, we performed immunofluorescent GFP staining on the ipsilateral DRGs of sham and MM bearing mice euthanized on post-surgical day 17 or 24 (Figure 6A, B). We found 5TGM1-GFP infiltration in the ipsilateral L2 DRG of all MM^D24^ mice confirming the results from the transcriptomic analysis (Figure 6C). Similar results were found in the ipsilateral L3 (data not shown). No GFP expression was detected in DRGs of MM^D17^ and sham controls (Figure 6B, C). Thus, our data indicate that MM has the capacity to metastasize to the peripheral nervous system, which occurs concomitantly to development of nociception.

**Figure 6.**
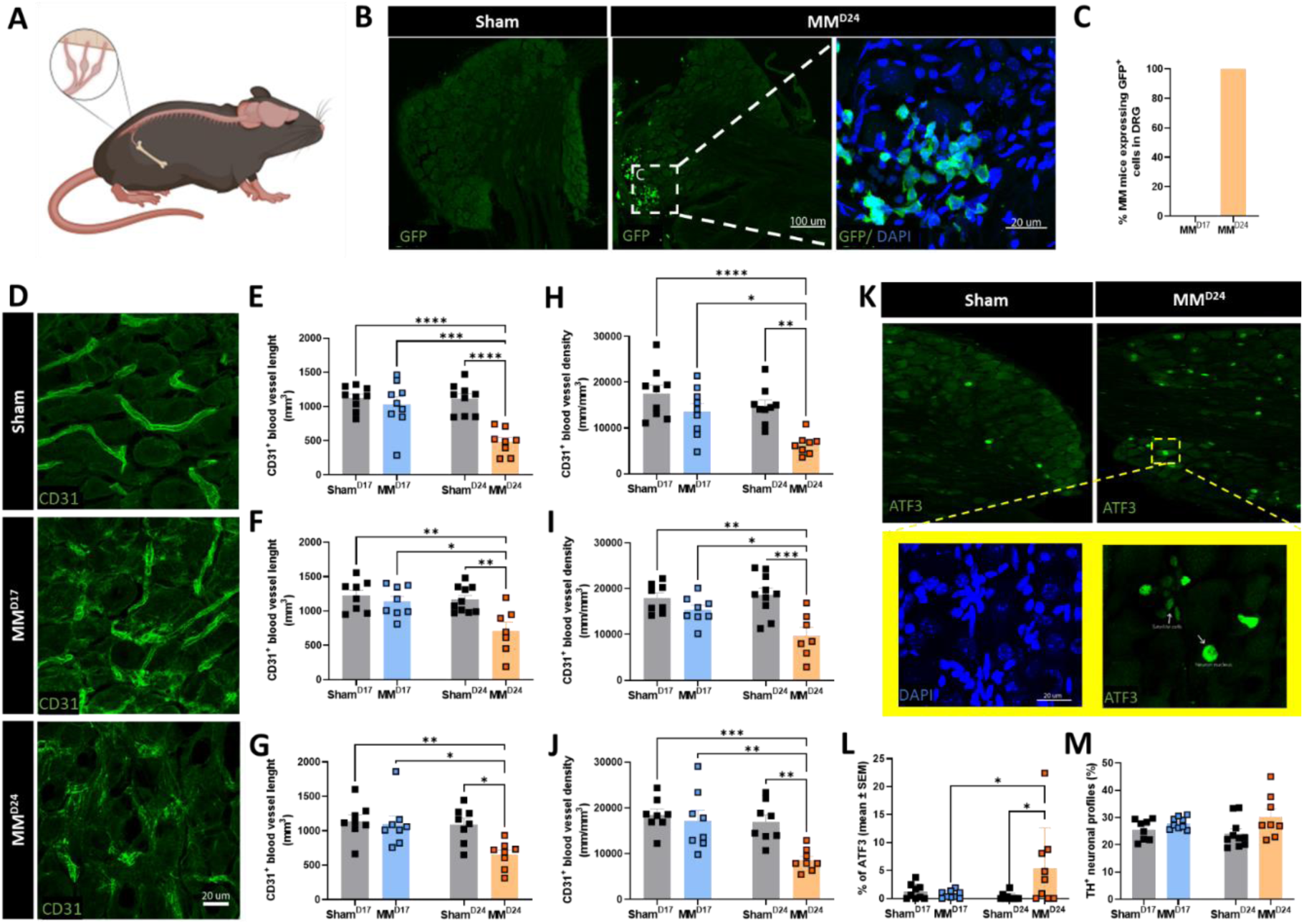
MM cells metastasize to the DRG causing damage to vasculature and neuronal bodies. (A) Ipsilateral and contralateral lumbar DRGs L2, L3 and L4 were collected. (B) Frozen sections from L3 DRGs were immunostained for GFP. (C) Number of MM bearing mice presenting GFP^+^ staining in the ipsilateral L3 DRG. (D) Representative images of CD31^+^ immunostaining in DRG frozen sections. Note the structural injury to CD31^+^ blood vessels in MM^D24^, indicated by white arrows. (E, F, G) Quantification of CD31^+^ blood vessel length in the ipsilateral L2 (E), L3 (F) and L4 (G) DRGs. (E) F(1,31)=10.57, p=0.0028, Sham^D24^ vs MM^D24^ p=0.0004; MM^D17^ vs MM^D24^ p<0.0001; sham^D17^ vs MM^D24^ p<0.0001; (F) F(1,29)=4.610, p=0.0403, Sham^D24^ vs MM^D24^ p=0.0105; MM^D17^ vs MM^D24^ p=0.0036; sham^D17^ vs MM^D24^ p=0.0017; (G) F(1,28)=4.321, p=0.0469, Sham^D24^ vs MM^D24^ p=0.0129; MM^D17^ vs MM^D24^ p=0.0142; sham^D17^ vs MM^D24^ p=0.0056 by two-way ANOVA followed by Tukeýs correction. (H, I, J) Quantification of relative CD31^+^ blood vessel density in the ipsilateral L2 (H), L3 (I) and L4 (J) DRGs. (H) F(1,31)=2.214, p=0.1468, Sham^D24^ vs MM^D24^ p=0.0119; MM^D17^ vs MM^D24^ p=0.0028; sham^D17^ vs MM^D24^ p<0.0001; (I) F(1,29)=5.178, p=0.0304, Sham^D24^ vs MM^D24^ p=0.0499; MM^D17^ vs MM^D24^ p=0.0005; sham^D17^ vs MM^D24^ p=0.0022; (J) F(1,28)=10.31, p=0.0033, Sham^D24^ vs MM^D24^ p=0.0037; MM^D17^ vs MM^D24^ p=0.0051; sham^D17^ vs MM^D24^ p=0.008 by two-way ANOVA followed by Tukeýs post-hoc test. (L) TH^+^ quantification in frozen sections from L2 DRGs. (K) ATF3^+^ immunoreactivity; exert denotes high-resolution imaging of ATF3^+^ and DAPI immunoreactivity on ipsilateral L2 MM^D24^ DRG. Note the presence of Nagoette nodules denoted with white arrows, suggestive of neuronal degeneration. (M) ATF3^+^ quantification in L3 DRGs. F(1,33)=5.363, P=0.0269; Sham^D24^ vs MM^D24^ p=0.0492; MM^D17^ vs MM^D24^ p=0.0239 by two-way ANOVA followed by Tukeýs post-hoc test. (O). MM= Multiple myeloma. Sham n=8-9; MM n= 7-10.

Next, we examined the integrity of DRG vascularization and potential neuronal damage through immunofluorescent staining of the ipsilateral DRG of sham and MM mice 17 or 24 days after cell inoculation. Our analyses of CD31^+^ blood vessels (Figure 6D) revealed a significant decrease in blood vessel length (Figure 6E-G) and density (Figure 6H-J) in DRGs from MM^D24^, but not MM^D17^, compared with sham at all the analysed lumbar levels. The proportion of sympathetic TH^+^ neurons was similar across samples (Figure 6 L); however, the percentage of activating transcription factor 3 (ATF3)^+^ neuron profiles was significantly increased in ipsilateral MM^D24^ as compared to sham^D24^, suggesting tumour-induced neuronal injury (Figure K, M). Additionally, ATF3 and 4′,6-diamidino-2-phenylindole (DAPI) staining revealed a specific pattern of nuclear staining consistent with the development of Nagoette nodules, indicative of neuronal degeneration (Peters et al., 2007) (Figure 6K). Finally, we examined the contralateral MM^D24^ DRGs and found intact vasculature and low levels of ATF3 expression (data not shown), suggesting that MM DRG infiltration and concomitant vasculature and neuronal damage may be a specific mechanism of MIBP.

### The transcriptomic DRG signature of a MM patient suggests metastatic infiltration

Following our unexpected observation of MM metastasis to the DRG of myeloma-bearing mice displaying MIBP, we questioned the translational validity of our findings. To evaluate whether the genetic signature of MM cells was present in peripheral nervous system of a MM patient, we accessed the transcriptome of 68 DRGs collected from 39 patients with 18 different types of cancer (Ray et al., 2022), as well as that of two thoracic DRGs from one MM patient. Median age of the patient cohort was 60 years (spamming from 33-79 years) and 15 patients were females (35.58%); detailed patient characteristics have previously been reported (Ray et al., 2022). First, we used publicly available datasets to generate a transcriptomic signature of composed of human markers generally expressed in MM cells (Zhan et al., 2006; Jang et al., 2019; Barwick et al., 2021) that show low or no expression in healthy DRG tissue (Ray et al., 2018). Datasets were chosen to represent different high throughput technologies (Affymetrix, bulk RNA-seq and single cell RNA-seq) and the final MM signature contained 40 genes that were ranked within the top 10,000 genes in Affymetrix and bulk-RNA, detected in over a third of single-cell RNA-seq cells, and had an expression rank <0.75 TPM in normal DRG tissue. Gene expression of the 40 signature markers was generally higher in the DRGs of the MM patient compared with most other cancer samples (Figure 7A). Indeed, the distribution of percentile gene expression of the two MM samples over the other cancer samples, across the 40 MM markers, showed a general shift towards higher percentiles (Figure 7B), with a mean percentile of 71,6 (mean of the two samples from the MM patient). In order to check whether other cancer samples displayed a similarly high or higher MM signature, we plotted the mean percentile marker distribution for all other 68 cancer samples and confirmed that MM samples showed among the highest mean percentile (Figure 7C). Other samples with a high mean percentile (indicative of similarly enriched expression for the selected marker genes) were DRGs from prostate and renal cell carcinoma patients (Supplementary File S2).

**Figure 7.**
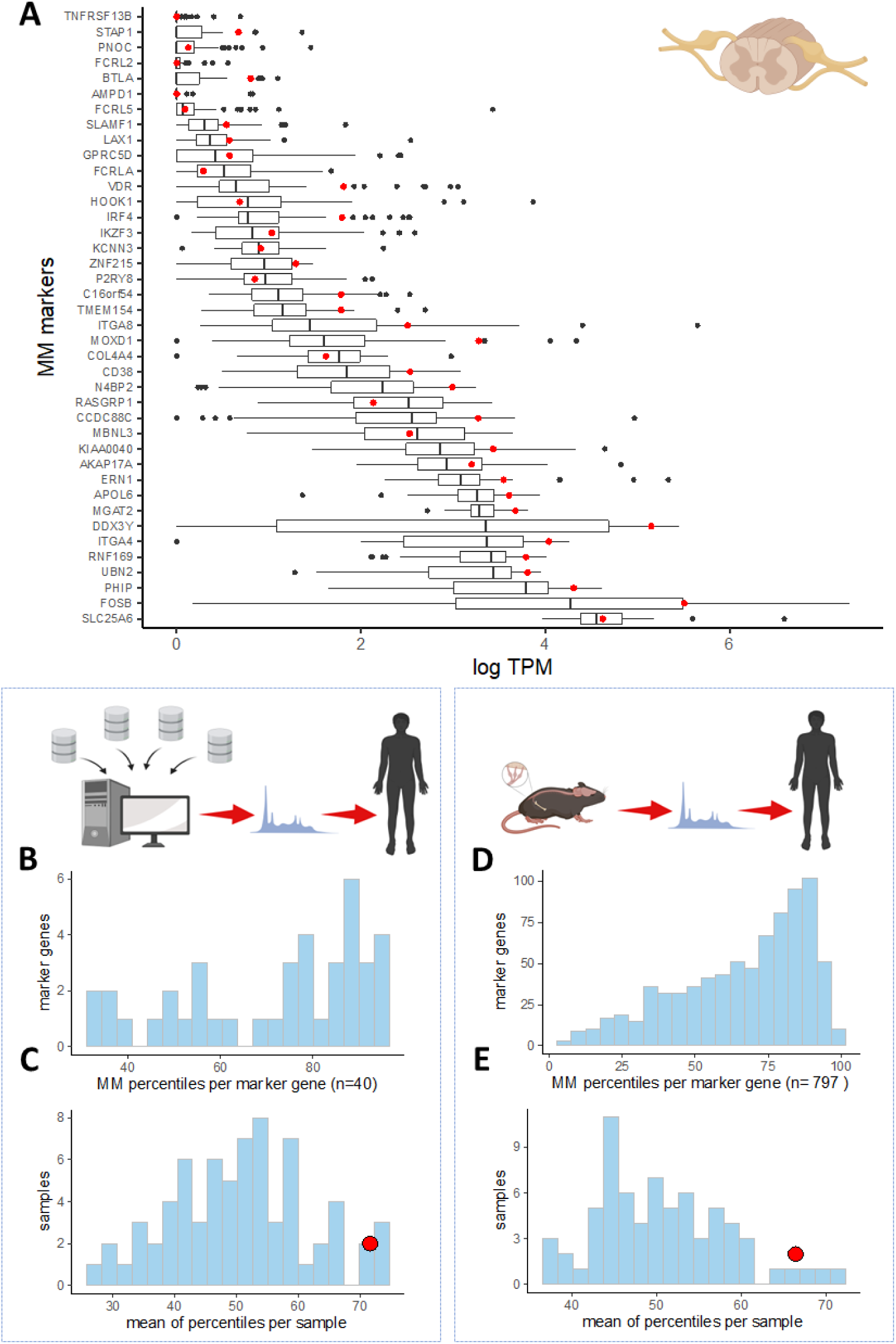
The transcriptomic signature of a MM patient suggest cancer infiltration to the DRG. (A) Box-plot of log FPKM counts for the 40 signature MM markers across 70 DRGs from cancer patients. Red dots indicate the mean expression level in two thoracic DRGs from one MM patient. (B) Distribution of percentile gene expression for all 40 markers in the DRGs of the MM patient. (C) Distribution of mean percentiles across all cancer samples; red dot indicates the mean percentile for the MM patient. (D) Distribution of percentile gene expression for the signature set derived from the mouse MM^D24^ upregulated DEGs (padj<0.05). (E) Distribution of mean percentiles across all cancer samples; red dot indicating the mean percentile for the MM patient, for the mouse MM^D24^ derived set.

Next, we asked if the MIBP transcriptomic signature identified in our animal experiments was translatable to the human condition. Since the DEGs observed in MM^D24^ vs sham^D24^ mice seemed to be largely driven by MM cell infiltration to the DRG, we hypothesized that these DEGs are either expressed in MM cells or disrupted as a result of MM infiltration and could be used as a proxy for MM metastasis in patient data. Thus, we selected the set of human orthologs to the mouse upregulated DEGs (padj<0.05) in MM^D24^, resulting in a set of 797 genes. Like with the human datasets, the relative gene expression of the chosen markers displayed a distribution shift towards higher percentiles, indicating a tendency towards overexpression of the genes from this set in the human MM DRGs compared to other cancers (Figure 7D). The MM samples showed among the highest mean percentile (mean of the two MM samples - 66,4) (Figure 7E). Taken altogether, our data suggest that cancer metastasis to the DRG may also occur in MM patients.

## Discussion

Bone pain remains among the main complaints from MM patients and significantly impairs their quality of life; indeed, previous reports have highlighted that MM patients report more symptoms and problems than leukaemia and lymphoma patients (Johnsen et al., 2009). However, the preclinical search for adequate analgesic options for MIBP is scarce and pathophysiological mechanisms underlying bone cancer pain are poorly understood (Hiasa et al., 2017; Olechnowicz et al., 2019; Diaz-delCastillo et al., 2020b).

In this study, we use our previously characterized local immunocompetent mouse model of MIBP (Diaz-delCastillo et al., 2020b) to investigate disease-driven alterations of the central and peripheral nervous system that may lead to rational search of new analgesic targets. Following 5TGM1-GFP cell transplantation into the intrafemoral marrow of a tumour-permissive mouse strain, we observed the progressive development of non-stimulus evoked nociceptive behaviours that can be considered as surrogate markers of spontaneous pain and/or wellbeing (Deacon, 2006; Sliepen et al., 2019). Behavioural tests and experimental time-points (i.e. collection of tissue on post-surgical day 17 or 24) were selected according to our previous model characterization (Diaz-delCastillo et al., 2019).

Indeed, we have previously shown that systemic opioid administration (10 mg/kg morphine) on post-surgical day 26 reverses the MM induced deficits in limb use, further confirming model validity (Diaz-delCastillo et al., 2019).

Following the direct transplantation of 5TGM1-GFP cells into a permissive microenvironment, we observed trabecular bone loss as early as post-surgical day 17, prior to the development of nociception. This apparent disconnection between osteolytic damage and nociception is consistent with preclinical and clinical evidence highlighting the limited analgesic efficacy of commonly used anti-resorptive treatments such as bisphosphonates (Mhaskar et al., 2017; Porta-Sales et al., 2017; Coluzzi et al., 2019; Diaz-delCastillo et al., 2019). In contrast, we observed cortical osteolysis at later stages of the disease, coinciding with the onset of nociception and suggesting periosteal involvement in MIBP. To confirm this, further immunostaining demonstrated 5TGM1-GFP cell escape to the periosteum of these bones, along with significant sprouting of sensory neurons, concomitant to nociception. Periosteal nerve sprouting has been posed as a potential mechanism of cancer-induced bone pain in animal models of breast (Bloom et al., 2011) and prostate bone metastasis (Jimenez-Andrade et al., 2010), as well as osteosarcoma (Mantyh et al., 2010), and anti-NGF therapy has recently showed modest results for the treatment of cancer pain (Sopata et al., 2015). Our results support the hypothesis that myeloma cells cause osteolytic cortical lesions through which they escape to the periosteum, where they may release neurotrophic factors that promote nerve sprouting and, potentially, nociception. The translational implication of these pre-clinical results is further supported by our exploratory observation of increased periosteal nerve density in NDMM patients compared to patients with hyperparathyroidism (Sayilekshmy et al., 2019), and the observation that periosteal nerve sprouting occurred more frequently in patients with periosteal infiltration of MM cells. Future studies addressing periosteal nerve sprouting in patients with MM and healthy controls are needed to confirm our findings.

To evaluate the contribution of periosteal nerve sprouting to MIBP, we tested the effect of repeated systemic NP137 administration. NP137 is a humanized monoclonal antibody targeting Netrin-1 that is currently undergoing Phase I and II clinical trials (NCT02977195; NCT04652076) as an anti-cancer treatment. Because Netrin-1 is a neurotrophic ligand involved in axon guidance, we expected that NP137 treatment would prevent periosteal nerve sprouting and, consequently, MIBP. Our results demonstrated that netrin-1 blockage effectively prevents periosteal nerve sprouting in myeloma-bearing mice without affecting bone morphometry. In contrast to previous studies (Fahed et al., 2022), systemic NP137 administration failed to have an effect of tumour burden or survival. While the reasons for this discrepancy are unknown, serum levels of sham drug-treated mice were twice as high compared to those of MM drug-treated mice, suggesting increased antibody clearance and volume distribution in MM-bearing mice. Future studies with greater doses of this antibody in mice with MM are warranted to determine its effect on tumour and disease progression.

Next, we evaluated the transcriptional DRG signature of MIBP and compared it to that of other pain models, including bone cancer metastasis, peripheral neuropathy, inflammation and chemotherapy-induced bone pain, as previously described (Bangash et al., 2018). Our data suggested that the transcriptional signature of MM bone pain was more similar to that of neuropathic and inflammatory pain than to the other pain models. While it has been speculated that central-acting agents targeting sensitization (i.e. gabapentin and antidepressants) may pose an alternative for the management of MIBP (Coluzzi et al., 2019), there is a lack of evidence supporting this treatment line. Detailed examination of the spinal cord in MM and sham mice revealed neither microglia activation not astrocyte reaction in the dorsal horn of the spinal cord at any investigated time-point.. However, central sensitization may occur without astrocytic or glial involvement, and we and others have previously reported that spinal microglial reaction in cancer-induced bone pain models is a highly variable occurrence that may not reflect the clinical reality (Honore et al., 2000; Diaz-delCastillo et al., 2020a). Instead, the neuropathic component of MIBP is in line with clinical reports describing baseline peripheral neuropathy in a proportion of MM patients (Richardson et al., 2012; Oortgiesen et al., 2022), which is also used as a prognostic factor of chemotherapy induced peripheral neuropathy and treatment outcome in this patient population (Dong et al., 2022). These results, in combination with our observation of periosteal nerve sprouting, suggest that medications targeted to neuropathic pain patients may be useful to treat a fraction of myeloma bone pain patients. However, our approach has several limitations, including that the GSEA comparison between our MM model and those described by Bangash et al (Bangash et al., 2018) is directly impacted by the quality of the data from each of the models. Thus, while the PSL model was the most similar to MM, it was also the one showing the highest statistical significance for the DEG analysis and resulted also in a higher number of significant GSEA terms; in contrast, cancer, CMP and MJP models (i.e. those with the lowest degree of similarity to MM) showed no DEGs falling within the threshold after FDR correction for the models, which may hinder the significance of our findings.

Among the most important findings from our mouse transcriptomics analyses was the high heterogeneity in the pattern of DEG in MM mice. Interestingly, the MIBP transcriptional signature was not correlated to surrogate markers of tumour burden or nociception but was instead highly dependent on the number of GFP counts. These results strongly suggest MM metastases to the ganglia, a never-before described feature of the disease that we further confirmed through immunohistological staining. Along with the spatial localization of MM cells in the ganglia, we observed structural damage and reductions in length and density of blood vessels innervating the lumbar ganglia. Whether damage to the blood vessels occurs prior to neoplastic infiltration (thus allowing the passage of MM cells into the DRG) or is a consequence of it, remains to be elucidated. In any case, neoplastic infiltration of the ganglia occurs concomitant to neuronal damage, as demonstrated by the increase in ATF3^+^ nerve profiles and the formation of nodules of Nagoette (Peters et al., 2007). This novel observation of neuronal degeneration at the onset of MIBP presents a new research avenue that requires further research to identify potential treatment targets; future studies could address the mechanisms of MM metastasis to the DRG and the involvement of immunomodulators in DRG colonization.

Our human transcriptomic analyses revealed a similar indication of possible neoplastic infiltration in the DRGs of a MM patient. To our knowledge, the only previous publication addressing this question is from 1958, when Dickenman et al. (Dickenman and Chason, 1958) found degenerative changes but no cancer infiltration in the DRGs of eight deceased MM patients. In contrast, we have performed bulk sequencing of two thoracic DRGs from a MM patient and compared our transcriptomic results to those of 39 patients with other cancer types. Comparing our custom-made MM transcriptional signature composed by MM genes with low or no expression in healthy DRG as per publicly available datasets to the gene expression profile of all 70 DRGs suggested that MM gene expression was enriched in the DRGs from the MM patient. Similarly, evaluating the expression pattern of human orthologs to DEG identified in MIBP mice revealed enrichment in the MM patient DRGs. This observational data supports the translational validity of our findings with the obvious limitation of the highly restricted sample number. Accessing quality human DRG tissue is challenging, but further research is needed to conclude whether cancer metastasis to the DRG is indeed a common occurrence in MM patients.

In conclusion, our data suggests that MIBP is mediated by concomitant mechanisms including periosteal nerve sprouting and neoplastic infiltration of the DRG. Moreover, the transcriptional signature of MIBP indicates a neuropathic component and pain management in MM patients may require a multi-targeted approach that include drugs targeting neuropathic pain.

## Acknowledgements

The authors thank Kaja Laursen, Camilla S. Dall and the animal technicians from the Department of Drug Design and Pharmacology at University of Copenhagen. We also thank the patients for providing their time and agreeing to be part of this project. Figures have been made in Biorender.

## Authorś contributions

MDC and AMH: study conceptualization, funding acquisition and manuscript preparation. MDC, LJ, MAL, TLA, JMJA and AMH: study design. MDC, TN, DMT, NACS, JAVM, LPG, HE, DN, PMD MAL, JMJA and AMH: mouse experiments and data processing. MDC, OP, LJ and AMH: planning, performing and analysing the transcriptomics mouse experiment. MDC, JC, TJP, PMD, OA: planning, performing and analysing human transcriptomics data. MDC, REA, AM, ADC, TLA and AMH: collection and analyses of human bone biopsies.

