## Supplementary 1 for "Metastatic infiltration of nervous tissue and periosteal nerve sprouting in multiple myeloma induced bone pain"

**SUPPLEMENTARY METHODS**

**Locomotor activity**

Distance and speed moved by freely walking mice was automatically measured with Ethovision XT 8.5 (Noldus Information Technology, Wageningen, The Netherlands). On post-surgical day 24, myeloma-bearing and sham mice were individually placed in standard, transparent plastic cages (125 x 266 x 185 mm) in a quiet, dimly illuminated room and their free movement was tracked in 1 min time-beans over a total of 15 minutes.

**Spinal cord tissue evaluation**

Spinal cords were serially sectioned at 30-µm thickness; lumbar regions L1 to L6 were identified as per their described grey matter anatomy (Watson et al., 2009) and collected. Slides were washed in PBS, permeabilized in 0,1% triton X in PBS and blocked with 2% BSA (Sigma Aldrich, Søborg, Denmark). Then, slides were o/n incubated with a rabbit antibody against Iba-1 (Wako Pure Chemical Industries, Ltd., Neuss, Germany), GFAP (DAKO Agilent, Glostrup, Denmark) or P-p38 (Thr180/Tyr182) (Bionordika, Cell Signaling Technology, Herlev, Denmark) at 4°C. Labelling was detected with an Alexa Fluor 594 goat anti-rabbit secondary antibody or Alexa Fluor 594 donkey anti-rabbit (1:300, Invitrogen, Themo Fisher Scientific, Slangerup Denmark). Slides were then washed, counterstained with DAPI (Themo Fisher Scientific, Slangerup Denmark) and mounted with Fluorescent Mounting Medium (DAKO Agilent, Glostrup, Denmark). Negative controls were performed by omission of primary antibody. Serial sections separated by at least 240 µm were imaged with a 20x objective using a Zeiss Axioskop 2 microscope with a high resolution Axiocam MRm camera (HAL100; Zeiss, Feldbach, Switzerland) and a fluorescent transmitter (HXP 120, Digital Scientific, Cambridge, UK). Pseudo-anonymised images were analysed in Image J (National Institutes of Health, NIH, Bethesda, USA) by a researcher blinded to experimental mouse and group. Briefly, positive Iba-1 and P-p38 cells were quantified within a pre-determined area of interest (347 x 260 µm^2^) located in the dorsal horn and including laminae I-IV. GFAP was quantified by measuring fluorescence intensity in the dorsal horn and calculating the corrected total cell fluorescence (CTCF) as previously described by us and others (Ansari et al., 2013; Diaz-delCastillo et al., 2018).

**Supplementary Table S1**

|  | Procedure description |
| --- | --- |
| Randomization | - Time-course experiment: mice were randomized into sham or MM according to their baseline burrowing performance and baseline weight. - Transcriptomics experiment: mice were randomized into sham or MM according to weight.   Randomization was performed by listing all mice from lower to higher baseline burrowing capacity or weight and stratifying in two groups. Average baseline burrowing capacity or weight was then calculated and mice were inter-exchanged among groups until the averages were equal ± 10%. |
| Allocation concealment | To minimize bias and cage, mice from different groups (i.e. MM and sham) were caged together. No physical distinctions among groups were available at mouse or cage level. The allocation code was kept in a close, secure file until the end of the experiment. |
| Blinding | All behavioural experiments were conducted by an investigator blinded to the experimental group. |
| Exclusion criteria | - In all experiments: mice displaying surgical complications. - In the time-course experiment: 10% worst burrowers (excluded only from burrowing behaviour). - In the transcriptomics experiment: mice with RNA RIN < 8.0 |
| Reporting of excluded animals | - Time-course experiment: - 2 mice due to surgical complications. - 10 mice due to cancer contamination in vehicle solution. - 2 mice in burrowing behaviour (10% worst burrowers) - Transcriptomics: 2 mice due to low quality RNA. |

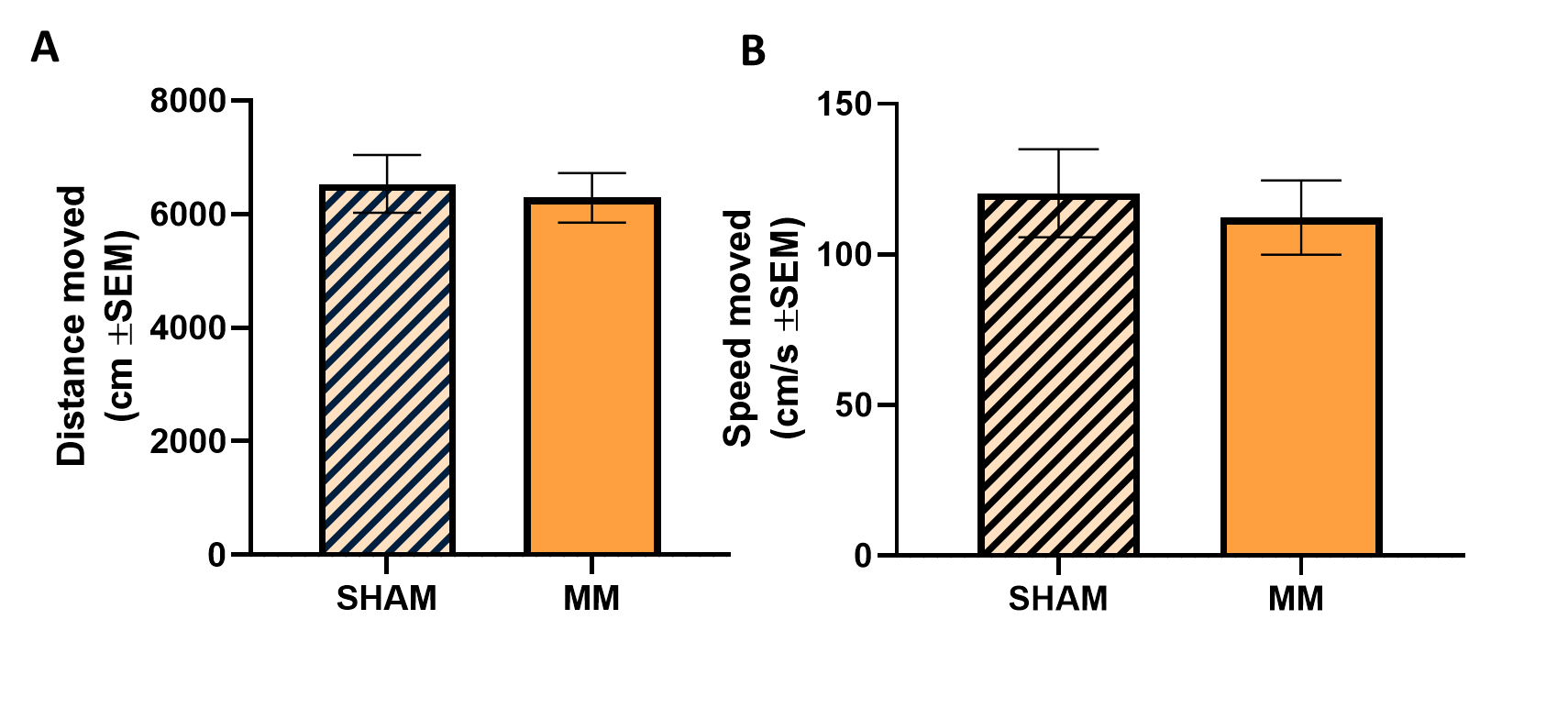

**Supplementary Figure S1. The mobility of myeloma-bearing mice was unaffected by disease progression.** On post-surgical day 26, distance moved (A) and speed moved (B) on MM mice and sham was not significantly different, indicating that the MIBP deficits in the limb use score were not cause by motor impairment. Data are presented as mean ± SEM. Sham n=4; MM n=10.

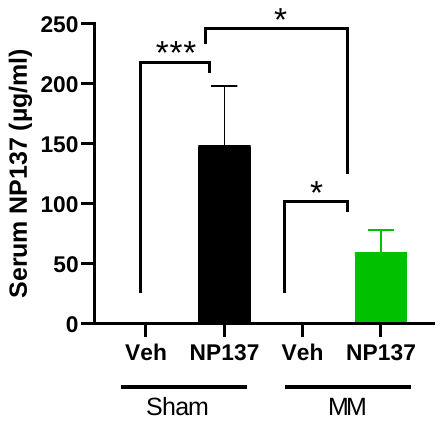

**Supplementary Figure S2. Serum levels of NP137.** The serum levels of sham drug-treated mice were higher than those of MM drug-treated mice; increased antibody clearance and volume distribution is often seen in cancer models, where antibody pharmacokinetics change in parallel to tumour size. Data are presented as mean ± SEM. Sham n=4; MM n=8/9.

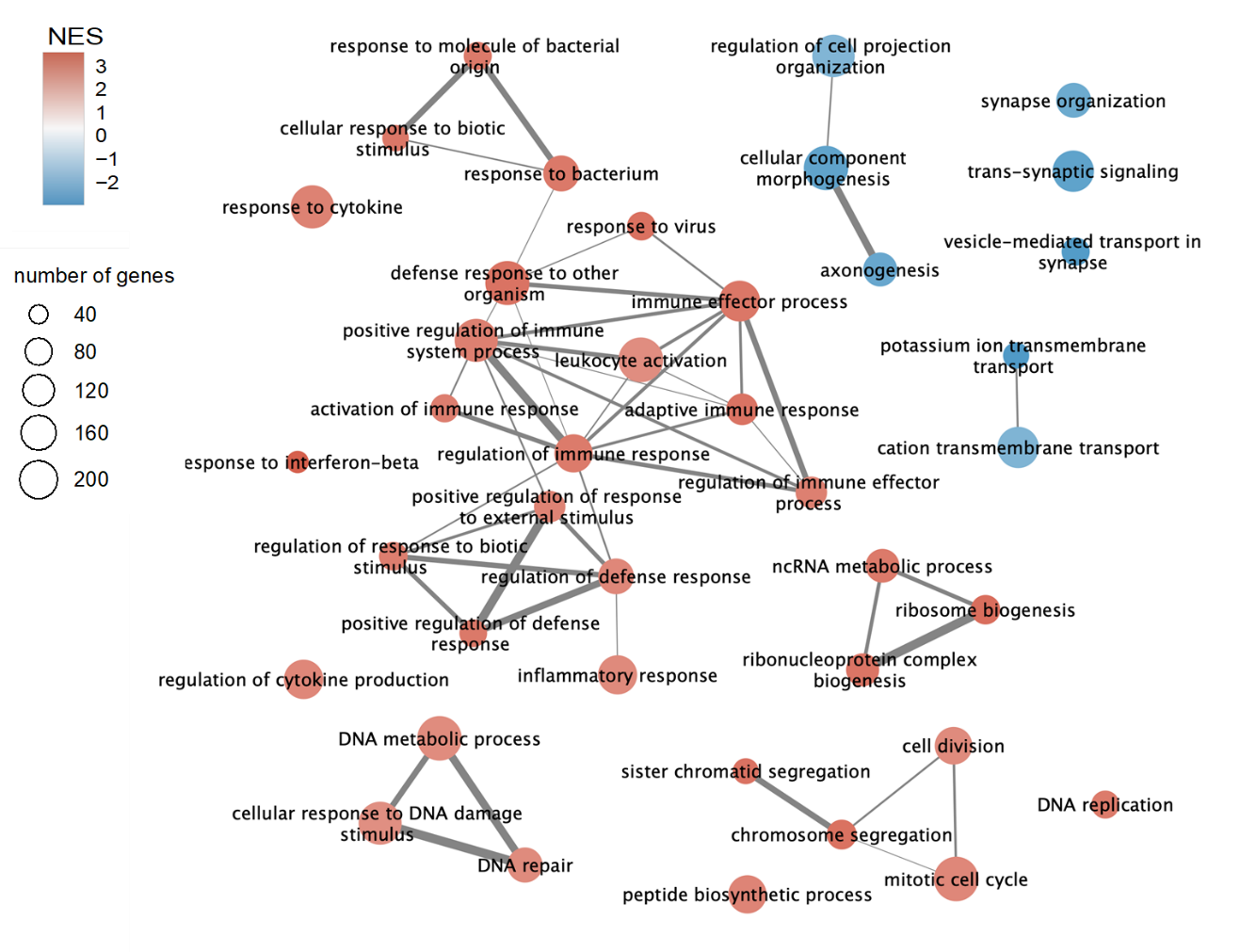

**Supplementary Figure S3. Top 40 GO BP terms from GSEA analyses.** Network view after filtering out highly similar terms. Node color represents the normalized enrichment score (NES) (red – positive, corresponding to upregulated terms and blue negative – downregulated terms). Node size is proportional to the number of genes associated to the term. Edges represent the JC between each two terms. Edge width is proportional with JC; only edges with JC>0.3 are shown. The main upregulated terms are related to immune response, cell cycle and DNA repair, while the downregulated terms are related to neuronal signaling.

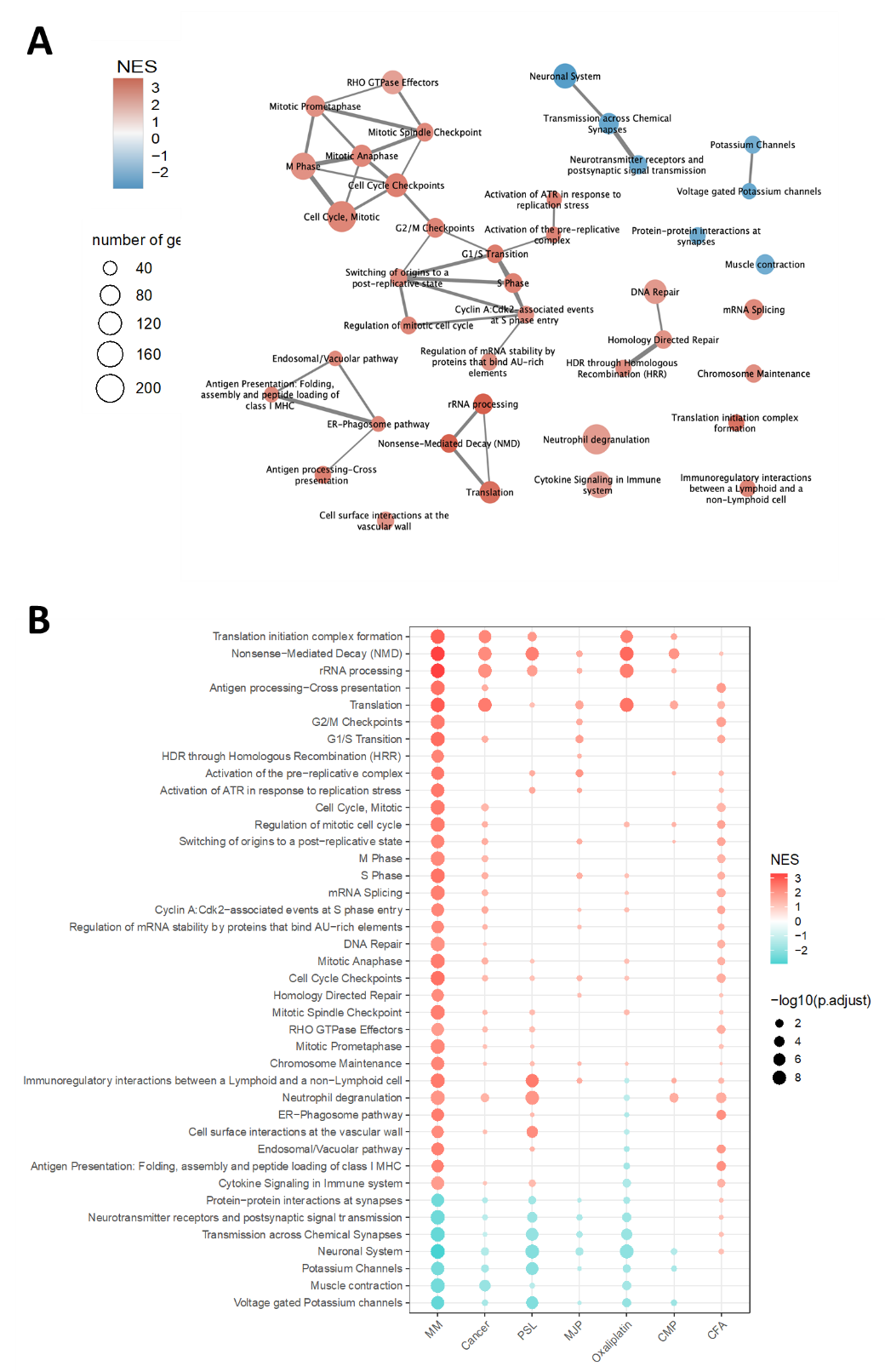

**Supplementary Figure S4.** **A**. **Network view of the top 40 enriched Reactome pathways from GSEA analysis.** Node color represents the normalized enrichment score (NES) (red – positive, corresponding to upregulated pathways and blue – negative, downregulated pathways). Node size is proportional to the number of genes associated to the term. Edges represent the JC between each two terms. Edge width is proportional with JC; only edges with JC>0.3 are shown. Most upregulated pathways are related to cell cycle, checkpoints, DNA repair, and downregulated pathways are related to neuronal signaling. **B.** Comparison of MIBP and the 6 models from Bangash et.al. in terms of GSEA for Reactome pathways. Normalized enrichment scores (NES) and corresponding adjusted p-values, for the top 40 enriched Reactome pathways in MIBP, are visualized across all seven pain models.

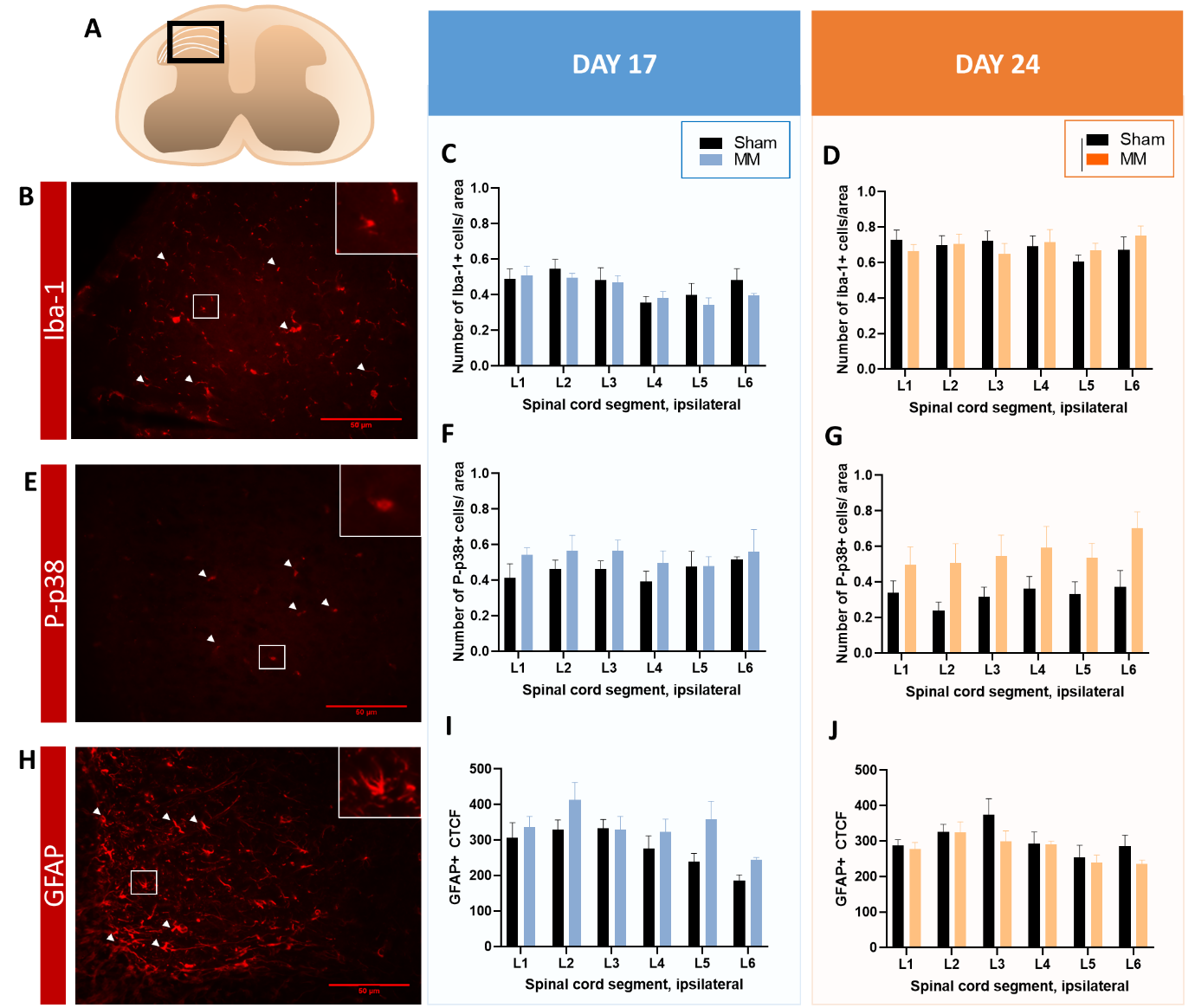

**Supplementary Figure S5. Microglia reaction and astrocytosis are not a main feature of MIBP.** (A) Overview of the 347 x 260 µm2 area encompassing laminae I-IV of the dorsal horn in which Iba-1, P-p38 and GFAP were analysed. (B, E, H) Representative image of Iba1^+^ microglia cells (B), Pp38^+^ microglia cells (E) or GFAP^+^ astrocytes (H) in the ipsilateral dorsal horn of the spinal cord (white arrowheads). (C, D, F, G) The relative number of Iba-1^+^ (C, D) or P-p38 (F, G) microglia cells in the ipsilateral dorsal horn of the spinal cord of lumbar regions L1 to L6 was unchanged between MM and sham mice euthanized on day 17 or 24. (I, J) The level of normalized GFAP expression in the dorsal horn of the spinal cord was similar between MM and sham mice euthanized on post-surgical day 17 or 24. L=lumbar. Data are presented as mean ± SEM. Sham n=6-8; MM n=7-8.
